# Increased pulmonary monocyte infiltration and attenuated phagocytosis defines perinatal dysfunction of innate immunity in Cystic Fibrosis

**DOI:** 10.1101/2023.09.11.556517

**Authors:** Florian Jaudas, Florian Bartenschlager, Bachuki Shashikadze, Gianluca Santamaria, Alexander Schnell, Nora Naumann-Bartsch, Heiko Bruns, Simon Y. Graeber, Andrea Bähr, Miquel Cambra-Bort, Michael Stirm, Jan Bernd Stöckl, Stefan Krebs, Helmut Blum, Christian Schulz, Dorota Zawada, Melanie Janda, Ignacio Caballero-Posadas, Karl Kunzelmann, Allessandra Morretti, Karl-Ludwig Laugwitz, Christian Kupatt, Armin Saalmüller, Thomas Fröhlich, Eckhard Wolf, Marcus A. Mall, Lars Mundhenk, Wilhelm Gerner, Nikolai Klymiuk

**Affiliations:** First Department of Medicine, Cardiology, Klinikum rechts der Isar, Technical University of Munich, School of Medicine and Health; Munich, Germany; Chair of Molecular Animal Breeding and Biotechnology, Gene Center, LMU Munich; Munich, Germany; Center of Innovative Medical Models (CiMM), LMU Munich; Munich, Germany; Institute of Veterinary Pathology, Freie Universität Berlin; Berlin, Germany; Laboratory for Functional Genome Analysis, Gene Center, LMU Munich; Munich, Germany; Department of Experimental and Clinical Medicine, University “Magna Graecia” of Catanzaro, Italy; Department for Pediatric and Adolescent Medicine, University Hospital Erlangen, Germany; Department of Internal Medicine 5, University Hospital Erlangen, Erlangen, Germany; Department of Pediatric Respiratory Medicine, Immunology and Critical Care Medicine and Cystic Fibrosis Center, Charité - Universitätsmedizin Berlin, corporate member of Freie Universität Berlin and Humboldt-Universität zu Berlin, Berlin, Germany; German Center for Lung Research (DZL), associated partner site, Berlin, Germany; Berlin Institute of Health at Charité – Universitätsmedizin Berlin, Berlin, Germany; Medizinische Klinik und Poliklinik I, LMU Munich, Germany; INRAE, Université de Tours, ISP, Nouzilly, France; Institut für Physiologie, Universität Regensburg; Regensburg, Germany; German Center for Cardiovascular Research (DZHK), Munich Heart Alliance; Munich, Germany; Institute of Immunology, University of Veterinary Medicine Vienna, Austria; present address: The Pirbright Institute, Woking, United Kingdom

## Abstract

In Cystic Fibrosis (CF) patients, cycles of infection and inflammation lead to fatal lung damage. While diminished mucus clearance is restored by highly effective CFTR modulator therapy, inflammation and infection persist in treated patients, suggesting alternative mechanisms may contribute to impaired immunity in CF. Here, we made use of a CF pig model to investigate the innate immune system at birth, before the onset of infection. We observed a substantial change in the composition of tissue resident immune cells towards emergency myelopoiesis, accompanied by increased infiltration of monocytes into CF lungs. A more immature status in the transcriptome profile of classical monocytes in CF pigs and preschool children with CF correlated with reduced phagocytic capacity, confirming a congenital and translationally conserved aberration of the immune system in CF. The lack of CFTR expression in circulating monocytes indicates an indirect etiology of these effects and suggests that additional immunological treatments are necessary for CF patients.

**One Sentence Summary:** Increased infiltration of lung tissue by monomyeloid cells and their impaired phagocytic potential cause dysfunctional imprinting of mucosal immunity in Cystic Fibrosis airways at birth and suggest early and specific treatment of the innate immune system in patients.

## INTRODUCTION

Cystic Fibrosis (CF) is the most common fatal autosomal recessive disease in Western countries, affecting multiple organ systems as a consequence of the defective ion channel Cystic Fibrosis Transmembrane Conductance Regulator (CFTR). The disease is characterized by a relentless cycle of gradually disintegrating host-environment interactions that destroys the airways and often causes premature death during the 3^rd^ or 4^th^ decade of life (Bell et al., 2020). At birth, airway structures appear grossly normal, but within the first months of life, the hallmark features of mucus accumulation, inflammation and infection manifest and lead to irreversible bronchiectasis (Esther et al., 2019; Rosenow et al., 2019). CFTR modulator therapies substantially slowed down the decline of lung function (Griese et al., 2021; Hisert et al., 2017; Middleton et al., 2019; Rowe et al., 2014), but airway deterioration seems unstoppable in older patients (Harris et al., 2020; Shteinberg and Taylor-Cousar, 2020). More recent clinical studies showed substantial residual airway inflammation and infection in patients treated with highly effective CFTR modulators (Casey et al., 2023; Nichols et al., 2023; Schaupp et al., 2023), pointing at the necessity to address the immune system in CF airways by alternative or additional therapies (Keown et al., 2020; Lara-Reyna et al., 2020). The detailed role of leukocytes in early CF pathogenesis remains poorly understood, in part because studies of blood samples from very young children are challenging (Hector et al., 2015) and systematic investigation of local immune cells in the airway wall is almost impossible (Regamey et al., 2012). Further, immune cells are highly sensitive to environmental cues (Lee et al., 2018). The imprinting of cellular stress pathways (Garg et al., 2012; Ribeiro and Boucher, 2010) or the priming of a pro-inflammatory milieu of CF patients (del Campo et al., 2011; O’Neill et al., 2018; Zhang et al., 2019) complicate mechanistic correlation of leukocyte deficits to primary CFTR dysfunction (Turton et al.).

In CF mice, standardized assessment of primary alterations in innate (Bonfield et al., 2012) and adaptive (Mueller et al., 2011) immunity has been performed. Interpretation of data, however, was hampered by the specific peculiarities of rodent innate immunity, ranging from different immune cell marker panels to distinct evolutionary shaping (Fairbairn et al., 2011; Flies, 2020). In addition, the influence of mucostasis on the immune system cannot be investigated in Cftr knock-out mice as they do not develop CF-like muco-obstructive lung disease (Grubb and Boucher, 1999; Rosen et al., 2018). Pig has been proposed as a better model species for human lung disease (Bertho and Meurens, 2021; Lunney et al., 2021; Rogers et al., 2008a) or inflammation (Meurens et al., 2012; Starbæk et al., 2018). Indeed, *CFTR^-/-^* pigs accurately reproduced malformations seen in early CF (Klymiuk et al., 2012; Rogers et al., 2008b). The neonatal CF pig revealed abnormal mucin production and reduced mucus velocity (Ermund et al., 2018; Hoegger et al., 2014) and less effective eradication of bacterial or viral pathogens (Berkebile et al., 2020; Pezzulo et al., 2012). Inflammatory markers were not upregulated in bronchoalveolar lavage (BAL) or lung tissue at birth (Bartlett et al., 2016; Stoltz et al., 2010). After a few days, inflammation triggered leukocyte infiltration into the interstitium and, even more pronounced, into the luminal space (Stoltz et al., 2010), similar to human patients (Regamey et al., 2012). A few studies included data on circulating leukocytes in the porcine CF model (Bréa et al., 2020; Gray et al., 2018; Paemka et al., 2017), but our understanding of lung immunity in early CF remains fragmentary.

To address the hypothesis that early changes in CF pre-sensitize the immune system for long-term deterioration, we aimed at comprehensive insight into neonatal leukocyte biology by characterizing major immune cell subpopulations in neonatal *CFTR^-/-^* piglets by flow cytometry, histology, transcriptome profiling and phagocytosis assays and comparative assessment of CF patients, including pre-school children.

## RESULTS

### Inflammatory status in CF airways and lung leukocytes

In line with previous reports (Bartlett et al., 2016; Klymiuk et al., 2012; Stoltz et al., 2010), larger bronchi of CF piglets showed thickening of the sub-epithelial smooth muscle layer at birth, but no further histomorphological lesions **(Fig. S1A)**. The weight of the respiratory tract and blood cell counts were not altered in CF piglets **(Fig. S1B, C)**. In proteomic data from bulk lung tissue **(Fig. 1A)** principal component analysis (PCA) or hierarchical clustering samples did not cluster by genotypes **(Fig. 1B, S2A)**. Based on the significantly altered proteins (25 of 3421 identified proteins), over-representation analysis (ORA) ranked “ferroptosis” and “immune response” from the KEGG and GOPB pathway database as over-represented biological processes **(Fig. 1C, D, Table S1)**. The most prominent hit, PAEP (fold change: 5.22; p<0.001) is the single ortholog of human PAEP/Glycodelin A, a verified stimulator of myeloid cells (Fig. S3A-E). Its abundance in the CF lung, however, likely arises from outside, i.e. from amniotic fluid (Halttunen et al., 2000), as *PAEP* transcription in the small airways was consistently low in WT and CF piglets **(Fig. 1E, S3G-I)**. Cytokine expression was comparable between genotypes at transcript level in airway tissue **(Fig. 1E)** and at protein level in BAL **(Fig. 1F)**. Similar to bulk tissue, airway-derived leukocyte proteome profiles were not separated by PCA or hierarchical clustering **(Fig. 1G, S2B)**, but ORA of differentially abundant proteins (120 of 3796 identified protein groups) pointed at a general activation of innate immune cells **(Fig. 1H)**. Strikingly, a number of differentially abundant proteins such as CD14 (log2fc:0.86; p=<0.001), ELANE (log2fc: 1.12; p=<0.001), MPO (log2fc:0.84; p<0.001) AZU1 (log2fc:0.86; p=<0.001), LTF (log2fc:0.97; p=<0.001) or ALOX15 (log2fc:0.89; p=<0.001) **(Fig. 1I, Table S2)** suggest a co-preparation of neutrophils with CF leukocytes or reflect the molecular signature of designated “neutrophil-like” or “granulocytic” monocyte populations (Ikeda et al., 2023; Yanez et al., 2017).

**Figure 1:**
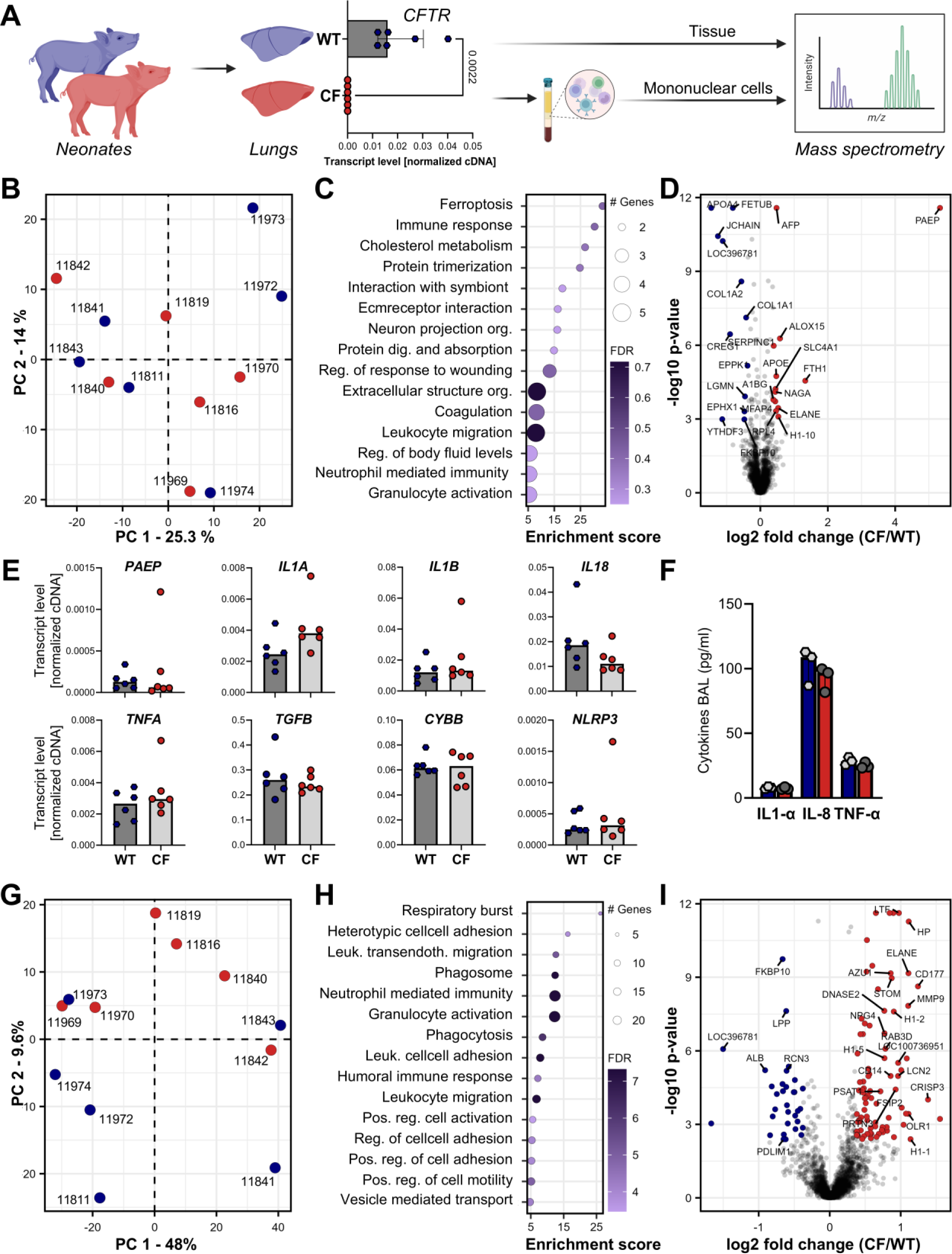
Proteome profile of neonatal CF lung tissue and its tissue-resident immune cells. (**A**) Bulk lung tissue and isolated tissue-derived mononuclear leukocytes of newborn CFTR^-/-^ piglets and littermate WT controls (n=6 each) were explored for proteomics. Unsupervised PCA (**B**), ORA **(C)** and differentially abundant proteins (**D**) in bulk tissue. LOC396781=IGHG. (**E**) Transcript levels of PAEP and cytokines in bulk tissue (n=6/genotype) as ΔCq values, normalized to GAPDH and ACTB (Fig. S3H, I). (**F**) Cytokine protein levels in BAL (µg/ml, n=3/genotype). Unsupervised PCA (**G**), ORA (**H**) and differentially abundant proteins (**I**) in isolated airway leukocytes. LOC100738951=diminished entry; LOC102160833=unclear; LOC100152327=IGLL5; LOC398781=IGHG. (C, D, H, I): Proteins with Benjamini–Hochberg (BH) corrected p-values ≤ 0.05 and fold-changes ≥ 1.5 were considered as significantly altered. (C, H): Top 15 gene sets according KEGG (Kyoto Encyclopedia of Genes and Genomes) and GOBP (Gene Ontology Biological Process). BH method was used for multiple testing adjustment. Dot sizes reflect the number of differentially abundant proteins (referred to as gene count in the figure) and color the significance of enrichment (FDR). Abbreviations: org.=organization; reg.=regulation; leuk.= leukocyte; pos.=positive. (D, I) Proteins in fold change vs. p-value. Red: significant upregulation in CF; blue: significant downregulation in CF. (E, F): Statistical analysis by Mann-Whitney U Test; none of the comparisons between genotypes in any assay reached significance level of p<0.05. Illustrations in Fig. 1.A created with Biorender.com.

### Elevated numbers of mononuclear phagocytes in neonatal CF airways

The absolute number of mononuclear cells (MNC) in blood or spleen were not altered between CF and WT **(Fig. 2A)**. In contrast, MNC were significantly increased in lungs (median: CF: 70.0 x10^6^ vs. WT: 38.5 x10^6^ cells per lung; p=0.007), while the absolute and relative weights of the respiratory tract appeared slightly reduced in CF **(Fig. S1B)**. Lymphocyte subsets in MNC, as determined by multicolor flow cytometry (FC) **(Fig. S4)**, were broadly similar in all three specimen. γδ T cells were substantially decreased in CF lungs (median: CF: 3.4% vs. WT: 7.7%; p=0.001) and CF blood (median: CF: 5.3% vs. WT: 10.6%; p=0.005) **(Fig. 2B, C)**, but subsequent analysis for their proliferative capacity **(Fig. S5A)** or separation into CD2^+^ and CD2^-^ subsets **(Fig. S5B)** did not indicate a mechanistic impetus. Similarly, the lower proportion of Ki-67^+^CD4^+^ T cells in the spleen **(Fig. S5A)** and the less differentiated status of circulating NK cells **(Fig. S5B)** in CF represented pronounced, but solitary findings.

**Figure 2:**
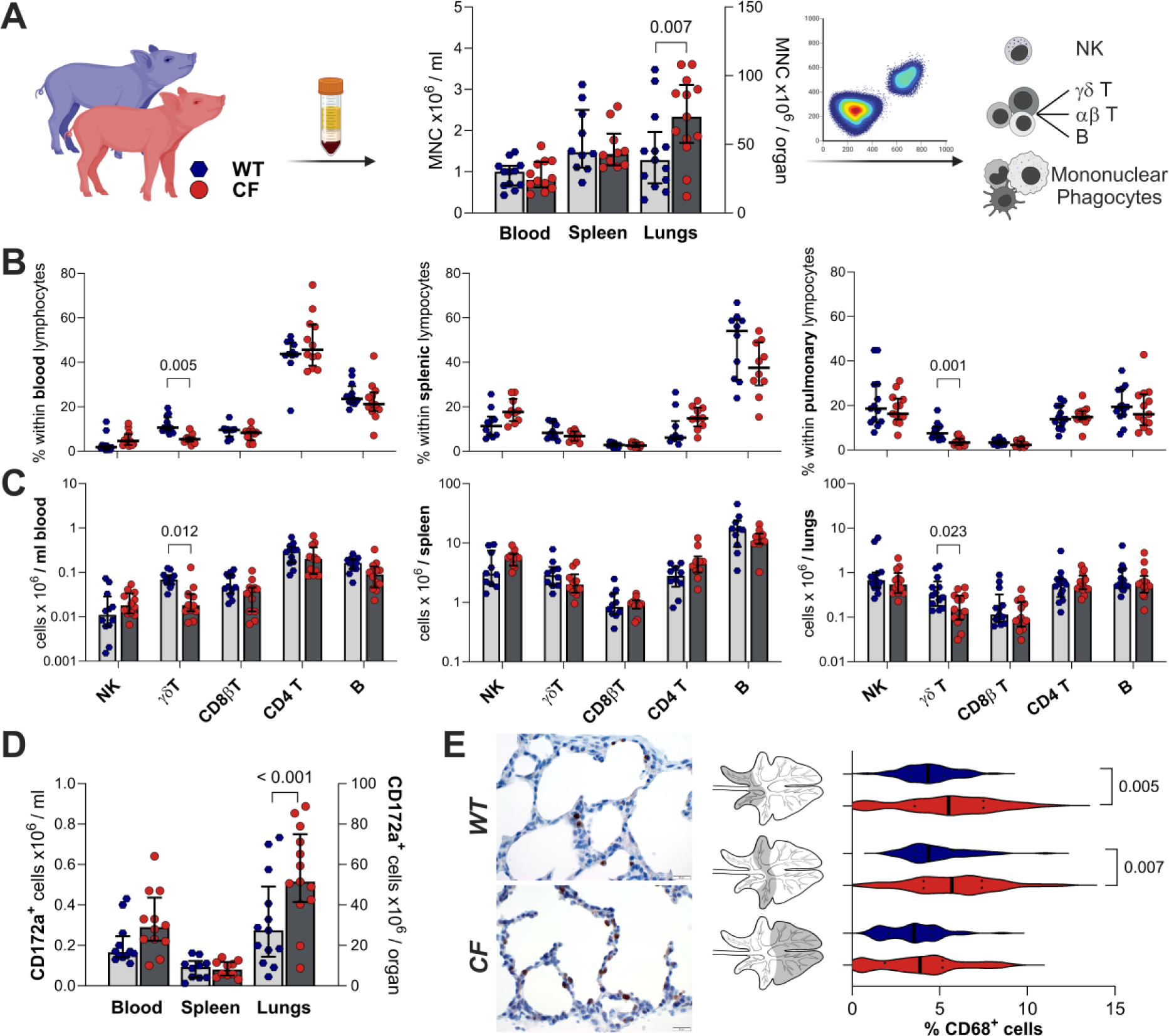
Immunophenotyping of major leukocyte subpopulations from CFTR^-/-^ piglets. (**A**) Mononuclear cells of newborn CFTR^-/-^ piglets and WT littermates were isolated by density gradient centrifugation from blood (n=12/genotype), entire spleen (n=10/genotype) or lung tissue (n=13/genotype). Leukocyte sub-populations were determined by FC (gating strategy in Fig. S4). Relative proportions (**B**) and absolute numbers (**C**) of major lymphocyte subpopulations. (**D)** Absolute numbers of CD172a^+^ MPs. A-D: Each data point represents one animal. Median ± IQR are shown. Significant Bonferroni-adjusted p-values are indicated, Wilcoxon matched pairs signed-rank test. (**E**) Immunohistological assessment of CD68^+^ cells in neonatal airways from CF (n=7) and WT (n=3) piglets. Left panel: Representative DAB-stainings with hemalaun counterstaining. Right panel: Quantification of CD68^+^ cells was performed by digital image analysis in 20 areas (2 x10^6^ μm^2^) per animal from cranial, cardiac and caudal lung lobes (middle panel). Percent CD68^+^ cells per section and genotype are expressed as median ± IQR and significant Bonferroni-adjusted p-values are indicated, Mann-Whitney U test. Illustrations in Fig. 2.A created with Biorender.com.

For their uncertain and divergent ontological origin (Guilliams et al., 2014), cells of the myeloid lineage were defined by a sequential gating strategy and expression of the porcine pan-myelomonocytic marker CD172a **(Fig. S4G)** and designated as mononuclear phagocytes (MPs). While no differences were found in blood or spleen, airway MPs were expanded in numbers (median: CF: 51.4 x10^6^ cells/lungs vs. WT: 27.5 x10^6^ cells/lungs; p<0.001) **(Fig. 2D)** and more cells had a mitotic phenotype (Ki-67^+^ median: CF: 55.5% vs. WT: 33.3%; p=0.01) **(Fig. S5A)**. Histological assessment with the macrophage marker CD68 in tissue sections identified MPs in the interstitium or in attachment to the alveolar wall of CF lungs **(Fig. 2E)**. Enrichment of CD68^+^ cells was specifically found in cranio-ventral compartments of apical (median: CF: 5.5% vs WT: 4.3%; p=0.005) and cardiac lobes (median: CF: 5.7% vs WT: 4.4%; p=0.007), but not in caudal regions of the diaphragmatic lobe. Changes in MP numbers and composition of airway MPs thus suggested investigation of their maturation processes.

### Altered plasticity of the perinatal respiratory network of mononuclear phagocytes

Alternative gating of isolated airway MNC with the pan-leukocyte marker CD45 and the pan-myelomonocytic marker CD172a **(Fig. S6A, B)** revealed non-hematopoietic CD45^-^CD172a^+^ cells (Dawson and Lunney, 2018) as a negligible fraction in the isolate and confirmed MPs as the dominant cell population **(Fig. 3A)**. Dimensionality reduction of polychromatic FC assays revealed a consistent pattern of CD172a, CD163 and MHCII expression in WT and CF within CD45^+^CD172a^+^ MPs **(Fig. 3B, 3C, S6C)**. CF-specific upregulation was observed for Ki-67 and CD14 **(Fig. 3D)**, matching the proteome profile of airway MPs **(Fig. 1I, Table S2)**. In perinatal WT lungs of pigs and humans, CD14^+^ MPs are commonly enriched (Maisonnasse et al., 2016) and diminish during differentiation into resident MPs thereafter **(Fig. 3E, S6D)**. In newborn CF piglets, the proportion of CD14^+^ MPs was outstandingly high (median: CF: 55.8% vs WT: 35.5%; p=0.026) and CD14 expression was increased (median MFI: CF: 1898 vs WT: 1207; p=0.004), while proliferation remained unchanged **(Fig. 3F)**. In absolute numbers, CD14^+^ MPs were more enriched (median: CF: 32.0 x10^6^ cells/lungs vs WT: 6.2 x10^6^ cells/lungs; p=0.004) than CD14^-^ MPs (median: CF: 24.4 x10^6^cells/lungs vs WT: 12.1 x10^6^ cells/lungs; p=0.026), but the latter showed higher proportions of Ki-67^+^ cells (CF: 47.3% vs WT: 33.0%; p=0.015).

**Figure 3:**
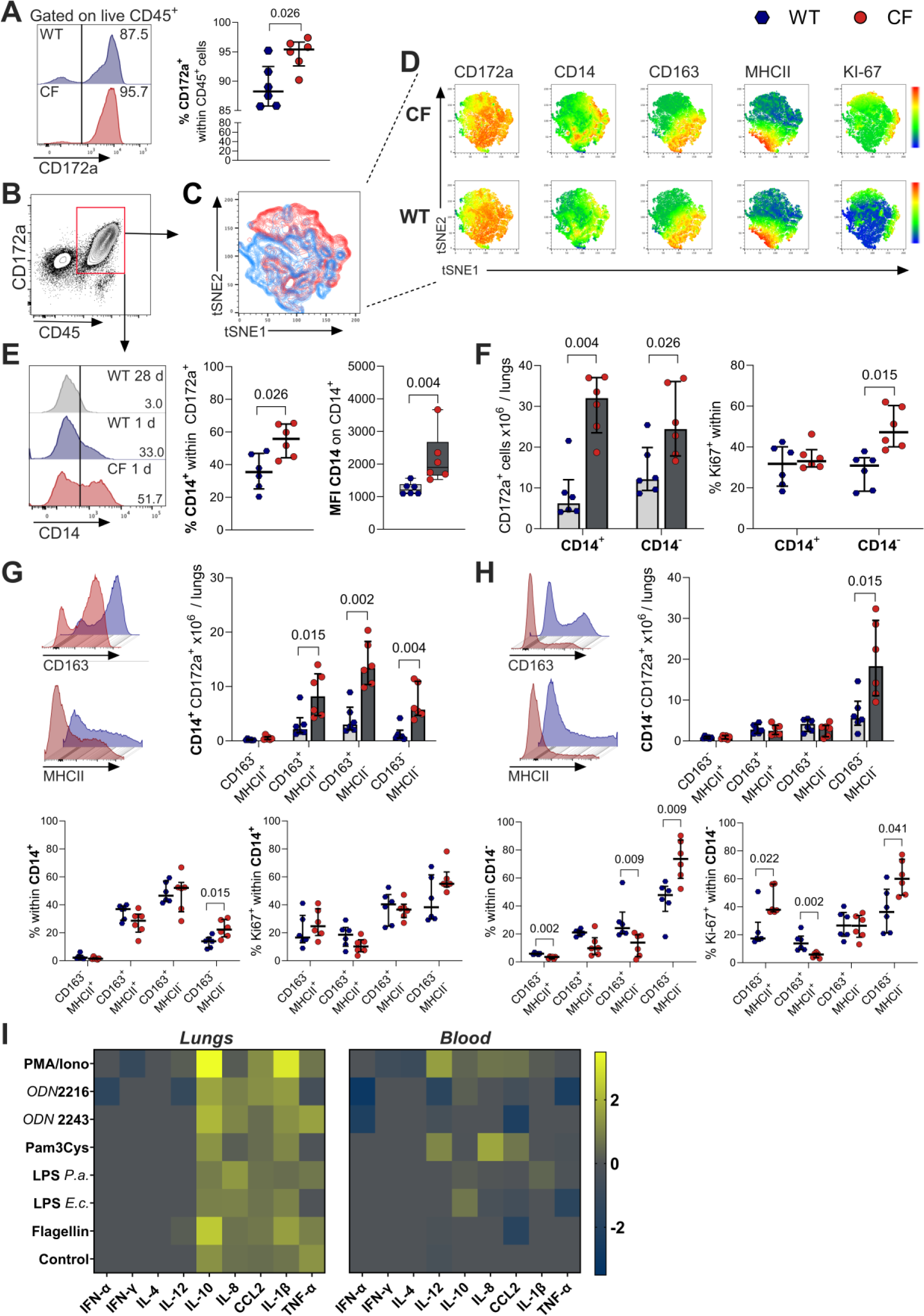
Altered plasticity of the CF respiratory Mononuclear Phagocyte (MP) Network. (**A**) CD172a expression on CD45^+^ pulmonary leukocytes. Left panel: representative histograms defining cut-off for CD172a expression. Right panel: Quantitative summary of CD45^+^CD172a^+^ MPs in CF (red) and WT (blue). (**B**) Gating of CD45^+^CD172a^+^ MPs for tSNE dimensionality reduction of multi-color FC (**C**). (CF=red, WT=blue). (**D)** tSNE plots separated for genotypes and selected markers. Expression levels indicated by color code (red=high, blue=low). (**E**) Left panel: Defining cut-off levels for CD14 expression in 28-days old WT. Middle Panel: Proportions of CD14^+^ MPs in CF (red) and WT (blue). Right panel: CD14 expression as mean fluorescence intensity (MFI). (**F**) MP subtyping by CD14 expression. Left Panel: Absolute numbers of CD14^+^ and CD14^-^ MPs per lung. Right Panel: Proportions of proliferating MP subsets, indicated as Ki-67^+^ cells. MP Subtyping with differentiation markers CD163 and MHCII in CD14^+^MPs (**G**) and CD14^-^ MPs (**H**). Upper left: Representative histograms for CD163 and MHCII. Absolute numbers (upper right), relative proportions (lower left) and proliferative capacity, indicated as Ki-67^+^ cells (lower right). (A-G) Each dot represents one individual animal (n=6 per respective genotype) and median ± IQR are shown. Significant p-values are indicated, Mann-Whitney U test. (**I)** Cytokine release from isolated leukocytes. Relative cytokine expression from CF and WT littermates (n=6 animals each), with log_2_ transformed ratios CF/WT indicated. Raw data in Figs. S7 & S8.

Subtyping with differentiation markers CD163 and MHCII (Maisonnasse et al., 2016) revealed an increase of all subsets within the CD14^+^ population in CF in absolute numbers, except for CD163^-^ MHCII^+^ cells **(Fig. 3G)**. In relative proportions, only the subset of CD163^-^MHCII^-^ MPs were enriched (median: CF: 22.3% vs WT: 14.0%; p=0.015) and proliferation appeared similar in all sub-types. Within CD14^-^ MPs, the CD163^-^MHCII^-^ population was expanded in absolute numbers (median: CF: 18.3 x10^6^ cells/lungs vs WT: 5.6 x10^6^ cells/lungs; p=0.015) and in relative proportions (median: CF: 87.2 % vs WT: 47.8 %; p=0.009), corresponding with a relative decrease in CD163^-^MHCII^+^ and CD163^+^MHCII^-^ cells **(Fig. 3H)**. Proliferation capacity of CD163/MHCII sub-types in CD14^-^ MPs appeared ambigous.

Pulmonary MNC from CF pigs revealed increased production of IL-10, IL-8, CCL2, IL-1β and TNF-α upon stimulation with PMA/Ionomycin, TLR agonists or even without stimulation **(Fig. 3I, S7)** while peripheral blood MNC (PBMC) did not indicate alterations in their stimulatory potential **(Fig. 3I, S8)**. Thus, we next aimed at deciphering the origin of airway MP alterations by investigating myeloid cells in the circulation.

### Disrupted monocyte homeostasis in peripheral blood

PBMC from newborn CF and WT piglets as well as from preschool children with CF and age-matched healthy controls (aged 2.4-5.4 years, **Table S3**) were pooled for each species-genotype constellation and analyzed for the transcriptome profile by single-cell RNA sequencing (scRNAseq) **(Fig 4A, S9A)**.

**Figure 4:**
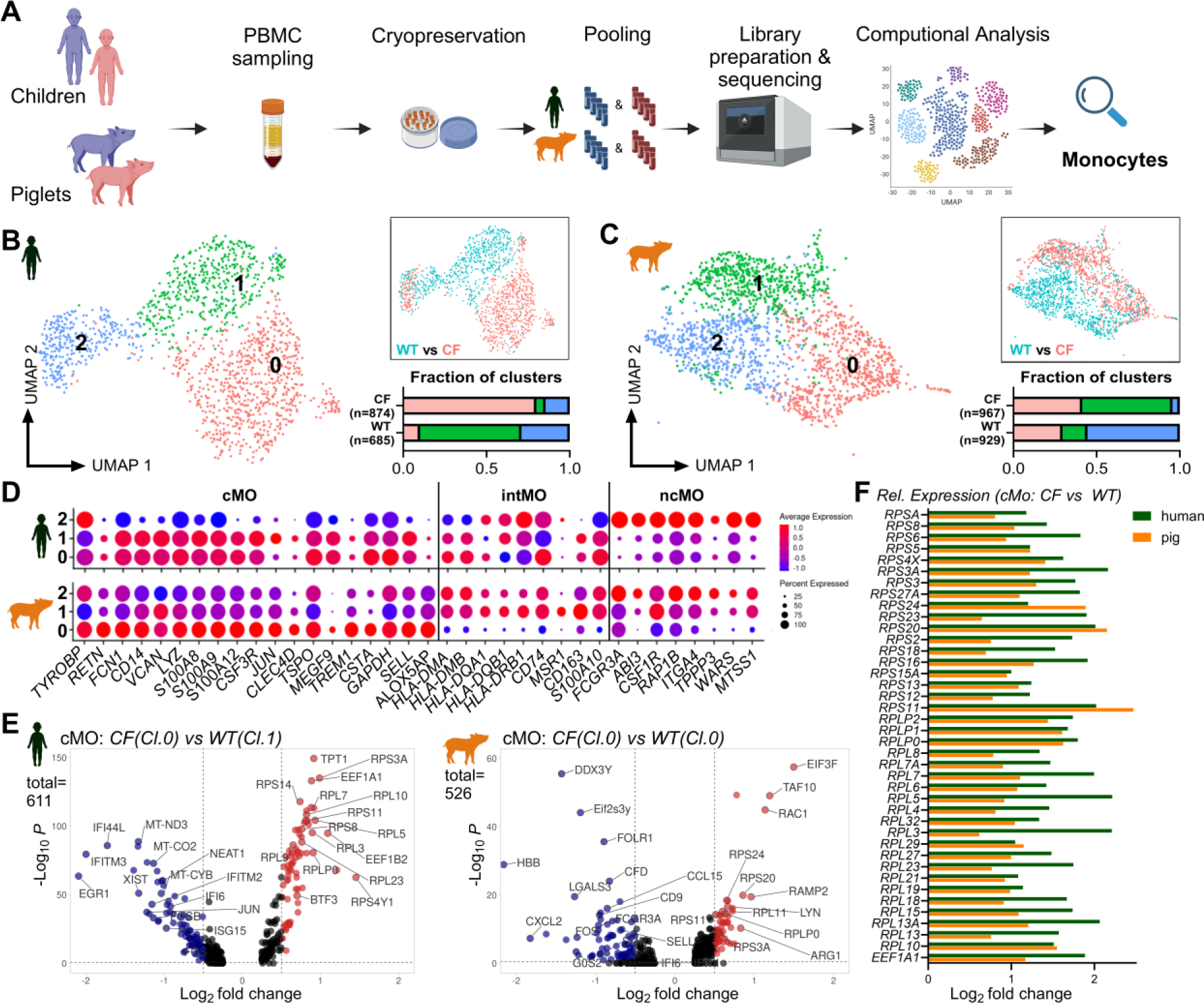
Disrupted monocyte homeostasis in peripheral blood during early CF. (**A**) Transcriptional single-cell profiling of circulating monocytes was done with pooled samples of purified PBMC from human infants and porcine neonates. Monocyte populations were identified as shown in Fig. S9. For human **(B)** and pig **(C)** monocytes, UMAP was used for unsupervised clustering (left panels) or allocation according to genotypes (upper right, red:CF, blue:WT). Proportions of each fraction in CF and WT are given (lower right, color according UMAP clusters). **(D)** Expression plots of selected markers for cMo, iMo, ncMo (see Tables S6 & S7) in human and pig. Dot size indicates the proportion cells expressing the respective marker within a cluster (“percent expressed”), color show the expression level of the markers (“average expression”). **(E)** Genes in cMo (human clusters 0 and 1, pig cluster 0) were considered as differentially regulated with –Log10 (p-values) and Log2 fold change <-0.58, >0.58. **(F)** Relative expression of ribosomal proteins (CF vs WT) in human (green) and pig (orange) cMo. Illustrations in Fig. 4.A created with Biorender.com.

Unbiased Uniform Manifold Approximation and Projection (UMAP) and association of data with a highly annotated scRNAseq reference of adult human PBMC (Bakken et al., 2021) confirmed clustering and identity of leukocyte sub-populations in human **(Fig. S9B, Table S4)**. In pig **(Fig. S9C)**, the pattern of established human immune cell markers appeared highly similar to the human data sets as well as in the literature **(Fig. S9D, E)**. Separation of B-, T- and NK-cells into sub-clusters was less pronounced in pig, confirming the primeval status of the adaptive immune system in newborns. Preschool children with CF indicated reduced naïve T-cell and enriched naïve B-and NK-cell populations **(Fig S9F, G)**.

Clusters 4 and 13 in human **(Fig. S9D)** and cluster 0 in pig **(Fig. S9E)** were designated as monocytes upon their marker pattern. UMAP of these populations revealed 3 defined sub-populations in both species, with clear separation according to genotypes **(Fig 4B, C, Table S5)**. Extended profiling with established markers of monocyte subpopulations **(Table S6, S7)** was done to allocate classical (cMo), intermediate (iMo) and non-classical (ncMo) monocytes **(Fig 4D)**. In human, both clusters 0 and 1 were designated cMo, while cluster 2 appeared as ncMo and iMo did not constitute, presumably due to low cell count and indifferent status (Kapellos et al., 2019). In pig, sub-populations 0, 1 and 2 rather reflected cMo, iMo and ncMo, respectively. In both, human and pig CF, ncMo (clusters 2) were reduced **(Fig. 4B, C)**. Differential expression analysis of cMo revealed a coherent pattern of gene encoding proteins of large (RPL) and small (RPS) ribosomal subunits **(Table S8)**. Both, preschool children and newborn pigs, showed consistently higher transcript levels of RPL and RPS in CF, compared to WT (approx. 2-fold) **(Fig. 4E, F)**.

### Attenuated phagocytic potential of circulating monocytes and granulocytes

Similar to bulk lung tissue and isolated airway MPs **(Fig. 1B, G, S2A, B)**, PBMC proteome profiles in the pig did not separate according to genotypes by hierarchical clustering **(Fig. 5A, S2C, Table S9)**. In line with airway MPs **(Fig. 1H, I)**, ORA and differential gene expression analysis of PBMC suggested changes towards neutrophils or neutrophil-like monocytes **(Fig. 5B, C)**, with specific lineage markers such as AZU1 (log2fc:0.76; p=<0.001), MPO (log2fc:0.55; p=0.028) or ELANE (log2fc:0.63; p=<0.001) being enriched. FC of CD172a^+^ MPs showed stable proportions of CD80/86^+^, CD163^+^ or CD16^+^ populations, while CD11b^+^ MPs were decreased in the circulation of CF pigs **(Fig. S10A-E)**. For CD80/86 and CD163, the expression level per cell remained unchanged in blood, spleen and lung **(Fig. 5D)**, whereas CD16 and CD11b abundance was diminished by half (MFI median blood: CF: 1130 vs WT: 2454; p<0.001; MFI median spleen: CF: 1866 vs WT: 2712; p<0.001; MFI median lungs: CF: 1453 vs WT: 2817; p<0.001) and 20% (MFI median blood: CF: 2458 vs. WT: 2951; p= 0.006; MFI median spleen: CF: 4090 vs WT: 5228; p=0.006; MFI median lungs: CF: 3943 vs WT: 5793; p=0.002), respectively. The vast majority of pig monocytes was able to ingest fluorescently labelled bacteria, but the uptake per cell diminished in CF (MFI median: CF: 673 vs WT: 1013; p<0.001) **(Fig. 5E, S10F)**. The decreased fraction of monocytes producing reactive oxygen species (ROS), (median: CF: 32% vs. WT: 63%; p<0.001) and the reduced amount of ROS per cell (MFI median: CF: 64.4 vs WT: 72.2; p=0.01) indicated impaired oxidative burst capacity in CF monocytes. Pig granulocytes ingested bacteria and produced ROS to a similar extent in WT and CF, but the phagocytic capacity per cell (MFI median: CF: 1058 vs WT: 1421; p=0.013) and ROS production per cell (MFI median: CF: 80.2 vs WT: 134; p=0.005) were significantly lower in CF pigs **(Fig. 5F, S10F)**. In patients with CF, reduced phagocytosis was was also found for *S. aureus* and *P. aeruginosa* in monocytes **(Fig. 5G)**. Accordingly, ROS production was reduced in human CF granulocytes after PMA stimulation **(Fig. 5H)**.

**Figure 5:**
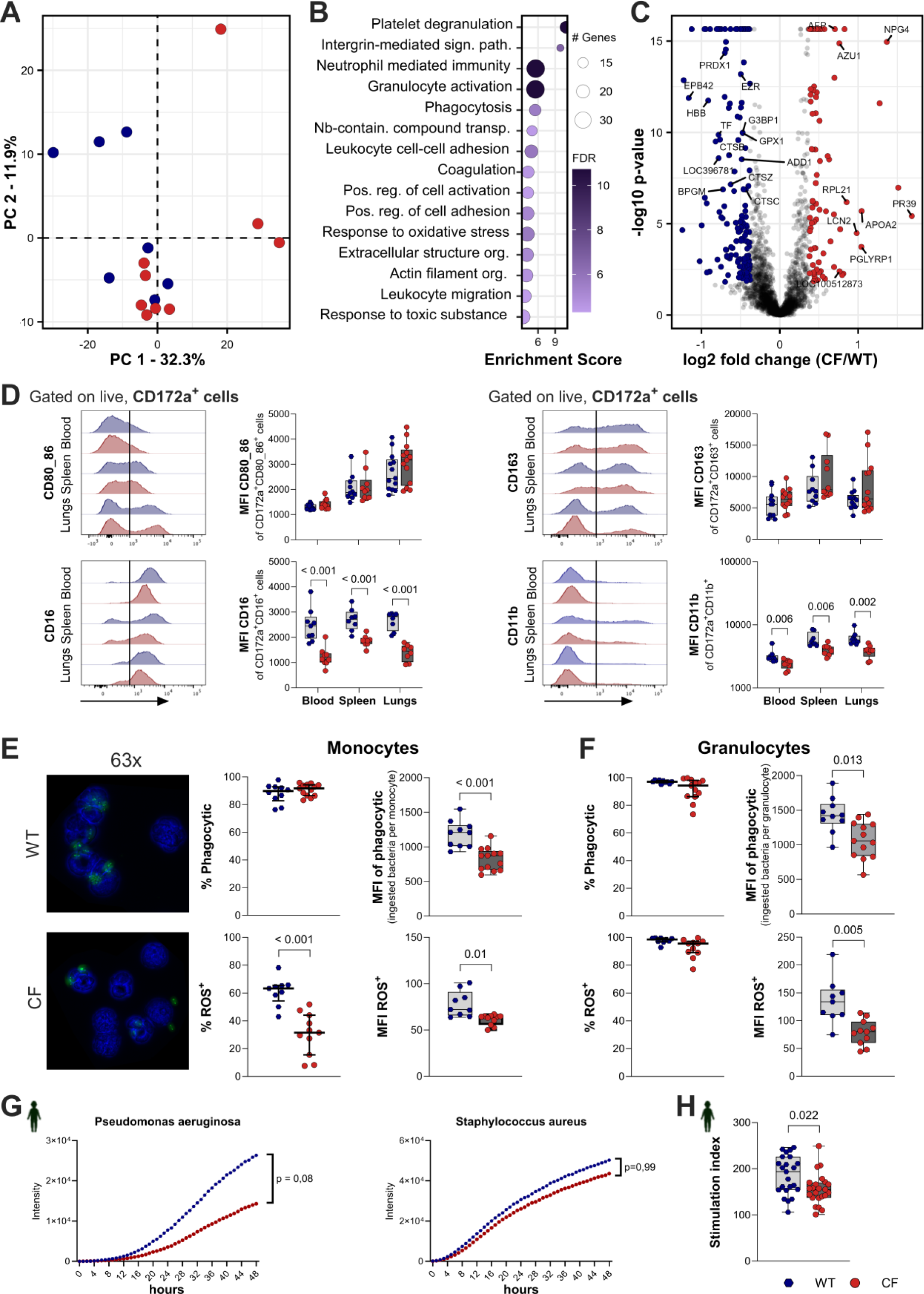
Attenuated phagocytic and oxidative potential of CF phagocytes. PCA (**A**), pathway enrichment analysis **(B),** and differentially regulated proteins (**C**) of proteome profiles in porcine PBMC, according to designations given in Fig. 1. Abbrevations in (B): sign.=signalling; path.=pathway; Nb= Nucleobase; contain.= containing; transp.=transport; pos.=positive; reg.=regulation; org.=organization. (C): LOC100154047=CTSG; LOC110256441=NPG3; LOC100512873= SLPI; LOC100152327=IGLL5. (**D**) Subtyping of CD172a^+^ MPs with CD80/86, CD163 (blood: n=12; spleen: n=10; lungs: n=13 per genotype), CD16 and CD11b (blood: n=9; spleen: n=8; lungs n=8 per genotype). Left Panels: Representative histograms indicating cut-off levels. Right panels: Marker expression, given as MFI per marker-positive cell. Phagocytic potential assessed for pig monocytes **(E)** and granulocytes **(F).** Quantitative evaluation was done for bacterial ingestion by detecting FITC-labelled E.coli (CF: n=13; WT: n=10) and for ROS production by detecting dihydro-rhodamine oxidation to rhodamine (CF: n=11; WT: n=9). Proportions of fluorescent cells and MFI per fluorescent cell were determined according to Fig. S10F. (E) Left panel: Representative staining of PBMCs ingesting FITC-labelled E.coli. green: FITC-label; blue: DAPI. (**H**) Uptake of pH-sensitive fluorescence-labelled S.aureus and P.aeruginosa by monocytes from human probands for 48 hours, as measured by fluorescence intensity. (**I**) ROS production in human probands after PMA stimulation is shown as stimulation index, indicated by MFI of stimulated and unstimulated cells. (D-H) Each symbol represents an individual animal; median ± IQR are shown. Significant Bonferroni-adjusted p-values are indicated, Mann-Whitney U test.

Over all, our data revealed enriched myelomonocytic cells in CF lungs already at birth with increased metabolic and mitotic activity, indicative of a more naïve state, while signs of inflammatory cytokine activation lacking. Genotype-specific signatures appeared also in PBMC of pre-school children and newborn pigs, correlating with impaired phagocytosis in circulating monocytes and granulocytes in both species.

## DISCUSSION

By investigating a pig model for CF and samples from CF patients, including preschool children, our study reveals that pronounced aberrations in the CF innate immune system are manifest already at birth and stretch out over years. The translational capacity of the pig model is demonstrated by concordant phagocytic impairment in CF myeloid cells in humans and pigs **(Fig. 5E-H)** and by the surprisingly coherent changes in human immune cell subsets on corresponding cell populations in the pig, specifically on monocytes **(Fig. 4D, S9D, E)**. Human markers from literature and databases **(Table S6, S7)** fitted even better to the pig cohort than to the human probands, demonstrating that differences between individuals were minimized by steady environmental conditions in animal housing and by comparing CF pigs with littermate controls to diminish genetic variability in the experimental cohort.

The findings of low and inconsistent numbers of leukocytes in BAL, the primeval status of the adaptive immune system **(Fig. S9C, E)** and lack of cytokine signaling **(Fig. 1E, F)** were in line with previous reports on newborn CF pigs (Bartlett et al., 2016; Paemka et al., 2017; Rogers et al., 2008b; Stoltz et al., 2010). The similar, if not lower, weight of distal airways **(Fig. S1B)** also matched other studies (Meyerholz et al., 2018). In contrast to the appearance of designated classical neutrophils in whole blood samples **(Fig 10F)**, such populations were completely removed in PBMC preparations by density gradient centrifugation **(Fig. S4A, S6B, C)**. Thus, we have a clear proof that the elevated number in airway leukocyte **(Fig. 2A, 3A)** represent respiratory MPs. Interestingly, increased numbers of MPs and their dominant localization to the interstitium **(Fig. 2E)** has been seen in human CF fetuses as well (Hubeau et al., 2001). The slightly higher stimulatory potential of isolated airway MPs **(Fig. 3I)** (Bruscia et al., 2009; Gray et al., 2018; Paemka et al., 2017) and the altered composition of airway MP sub-types **(Fig. 3F-H)** may therefore constitute already at a prenatal stage. Commonly, tissue colonisation by fetal monocytes is characteristic for mammalian species (Guilliams and Svedberg, 2021), but the numbers of CD14^+^ airway MPs, their relative proportions as well as their CD14 expression levels **(Fig. 3E)** in CF pigs were outstanding and match the dominant appearance of naïve CD14^+^MHCII^-^ cells in airway MPs of human pre-term infants (Prince et al., 2014). In further analogy to CF, a hyper-inflammatory milieu (Eldredge et al., 2019) establish as a long-term consequence of bronchopulmonary dysplasia in these patients.

While the increase in CD14^-^ MPs in CF may have derived from proliferation of tissue-resident progenitors **(Fig. 3F)**, the rise in non-proliferating CD14^+^ MPs in lung and the consistent enrichment of granule proteins in airway leukocytes **(Fig. 1I)** and blood **(Fig 5C)** clearly point at an increased infiltration of monocytes. According to previous work (Khmaladze et al., 2014), the diminished ROS production by CF MPs **(Fig. 5E)** may have directly caused the drop in γδ T cells in blood and lungs **(Fig 2B, C)**, a cell population that normally appears early during fetal development in mucosal tissue (Khairallah et al., 2018; Shibata, 2012). Upon airway infections, γδ T cells have been found to control inflammation (Kirby et al., 2007) and their lack was correlated to hyperinflammation in previous studies (Omar et al., 2020; Wehrmann et al., 2016). The proposed role of γδ T-cells as as a bridge or intermediate between innate and adaptive immunity (Ferreira, 2013; Hayday, 2019; Vantourout and Hayday, 2013) provides a mechanistic link between the alterations in myeloid cells and changes in the adaptive immune system in early CF **(Fig. S9F, G)** (Hausler et al., 2002).

The diminished bacterial uptake and ROS production in monocytes and neutrophils of newborn CF pigs **(Fig. 5E, F)** demonstrate that dysfunction of phagocytes is an early, if not confounding effect in CF. Both processes can be seen in correlation to reduced expression of CD16 or CD11b on CF MP in blood, spleen and lung **(Fig. 5D)** (Simonin-Le Jeune et al., 2013), which may reflect the shift towards a more naïve monocyte population, with reduction in ncMo **(Fig 4B, C)**. The recapitulation of the transcriptional changes in monocytes of CF pigs in preschool children with CF **(Fig. 4B-F, S9B, C)** as well as the reduced phagocytic potential and ROS production in CF patients of various ages **(Fig. 5G, H)** demonstrate that peri-natal changes in CF immunity have long-term consequences and fit the concept of “imprinted” or “trained” innate immunity (Hajishengallis et al., 2019; Saeed et al., 2014). Specifically, the changes in interstitium-resident MPs numbers and composition at birth stimulate speculations about consequences for the activity of their progeny alveolar macrophages (Guilliams et al., 2013; Tan and Krasnow, 2016) later on. Absence of common neutrophils in PBMC FC **(Fig. S4A, S6B, C)** and the lack of a designated neutrophil cluster in PBMC scRNAseq **(Fig. S9B, C)** were in stark contrast to the abundance of granule-related proteins in airway leukocytes **(Fig. 1I)** and PBMC **(Fig. 5C)**. In the literature, similar findings were explained by the presence of low-density neutrophil (Carmona-Rivera and Kaplan, 2023; Hardisty et al., 2021) or neutrophil-like monocyte (Ikeda et al., 2023; Yanez et al., 2017) populations. Both, if there was a difference between them, were primarily associated with non-homeostatic conditions and their dissection from monocyte lineages seems difficult (Wolf et al., 2019). At protein level, granule constituents appear early and remain throughout the neutrophil-lineage whereas transcription seems downregulated during their maturation (Grassi et al., 2018), reflecting to the enrichment of granule proteins in CF, while transcript levels in monocytes do not appear regulated **(Table S10)**. The maturation process of myeloid cells is also characterized by the downregulation of ribosomal activity (Hoogendijk et al., 2019), **(Table S8)**. Increased levels of ribosomal proteins in CF cMo from human and pig **(Fig. 4E, F)** suggest a more immature status, being in line with a proposedly altered maturation, a more naïve myeloid population or “emergency myelopoiesis” in CF at perinatal stage.

The perinatal changes in CF immunity have not yet been conclusively linked to the lack of CFTR expression in immune cells. Studies have shown at least some improvement in myeloid function in patients treated with CFTR modulator therapy (Cavinato et al., 2023; Zhang et al., 2023) or suggested a therapeutic capacity of WT hematopoietic stem cells in immune-depleted CF mice (Brinkert et al., 2021). Previous data have suggested CFTR transcripts in diverse granulocyte, monocyte or lymphocyte lineages (Yoshimura et al., 1991), albeit at a much lower level than in epithelial cells. Although we cannot rule out expression in myeloid cells such as airway or monocyte-derived MPs, there is strong evidence that *CFTR* was not expressed in the monocyte populations that we explored in scRNAseq. Data in pig (>200,000 reads/cell) and human (>50,000 reads/cell) indicated a similar number of expressed genes in PBMC (approx. 1500), but failed to detect any *CFTR* transcripts, rejecting a direct relationship between depleted CFTR function and impaired phagocytosis in circulating monocytes **(Fig. 5E)**. Considering the dynamic regulation of *CFTR* during embryonic development (Gaillard et al., 1994; Tizzano et al., 1994), it cannot be excluded that CFTR is shaping the immune system by expression in early progenitors. Alternatively, indirect pathogenic effects such as altered mucus processing and mucociliary clearance in early CF (Ermund et al., 2018) may enforce infiltration of monocytes into the CF lung. At least in a muco-static mouse model that is driven by Scnnb1-overexpression in the airways, non-CFTR provoked immune response were described (Hey et al., 2021) and correlated to hypoxia (Fritzsching et al., 2015). Very recently, upregulation of alternative ion channels, including SCNN1B, hypoxic responses and inflammation appeared as a general property of muco-obstructive airway diseases (Mikami et al., 2023). The restoration of mucus transport but incomplete correction of the immune system in CF patients treated with CFTR modulators (Casey et al., 2023; Nichols et al., 2023; Schaupp et al., 2023), however, does not support the hypothesis of mucus disruption and consequent infections as the only cause of chronic airway inflammation in CF.

Overall, our data suggest that aberrations in the CF immune system appear already at birth and constitutes early and long-lasting effects such as lung infiltration by MPs, their increased cytokine stimulatory potential, changes in circulating monocyte population including impaired phagocytic capacity and a reduced proportion of γδ T-cells. As a consequence of a pre-disposed hyperinflammatory immune status and improper response to bacterial challenges in CF, treatment strategies in patients should specifically include a comprehensive restoration of the innate immune system at an early stage of life.

## MATERIALS AND METHODS

### Regulatory statement

All work on CF pigs has been conducted under the supervision of the responsible regulatory authority (Regierung von Oberbayern) and animals were sacrificed for scientific purposes according to the German Animal Welfare Act. Blood sampling from preschool children with CF and age-matched healthy controls has been approved by the ethics committee of the Charité - Universitätsmedizin Berlin (EA2/016/18) and written informed consent was provided by the parents or legal guardians.

### Sampling

Experimental animals were produced from a *CFTR^+^*^/-^ breeding cohort (Ermund et al., 2018; Giorgetti et al., 2021; Klymiuk et al., 2012). *CFTR*^-/-^ null and *CFTR*^+/+^ wild type piglets were genotyped and sacrificed within the first day of life. MNC were immediately isolated from pig lung and spleen as well as pig or human blood by gradient centrifugation and stored at -80°C for protein analysis and in liquid nitrogen for FC. For molecular analysis, lung tissue and spleen was shock frozen on dry ice and stored on -80°C. For histopathological analysis, samples were immersion fixed in 4% formalin.

### Proteomics

Frozen lung tissue samples were powderized under liquid nitrogen and stored at -80°C. Cryopreserved leukocytes from airways and blood were directly used. Nano-liquid chromatography (LC)–tandem mass spectrometry (MS) analysis was conducted as previously described (Stirm et al., 2021). In brief, samples were lysed in urea / ammonium carbonate and enzymatically digested. Mass spectrometry was done on an UltiMate 3000 nano-LC (Thermo Fisher Scientific, Waltham, MA, USA) system coupled online to a Q Exactive HF-X instrument (Thermo Fisher Scientific). Analysis was done with MaxQuant (https://www.maxquant.org/, v.1.6.7.0) using Sscrofa as reference for protein identification. Principal component analysis and hierarchical clustering were used for unsupervised clustering. Over-representation analysis for significantly altered proteins was performed in WebGestalt (Liao et al., 2019).

### Quantitative reverse transcription PCR

RNA was isolated from powderized lung tissue by Rneasy Micro (Qiagen, Hilden, Germany) and cDNA was synthesized with SuperScript III (Thermo Fisher Scientific). RT-PCRs were run with HotStarTaq DNA polymerase (Quiagen). Quantification was done by qPCR using 2.5 µl cDNA, FastStart Essential DNA Green Master (Roche Life Science, Basel, Switzerland) in a LightCycler96 RT PCR system (Roche Life Science). *ACTB* and *GAPDH* were utilized for normalization.

### Immunophenotyping

Cryopreserved leukocytes were thawed, washed and stained. 5 x10^4^ – 2 x10^5^ leukocytes were examined per sample with a FACSCanto II (BD Biosciences, Franklin Lakes, NJ, USA), according to established multi-color protocols (Cossarizza et al., 2019). Data were analyzed with FlowJo™ software version 10.4.2 (BD Biosciences). Dimensionality reduction was done by tSNE.

### Histopathology

Fixed tissue was embedded in paraffin and stained with HE or a polyclonal anti-CD68 antibody (abcam, Cambridge, UK) that was visualized by a biotinylated goat anti-rabbit antibody (BA-1000; Vector Laboratories, Newark, CA, USA) and a avidin – horse radish peroxidase (HRP) conjugate (Vector Laboratories)Sections were counterstained with hemalaun and explored by digital image analyses (Dietert et al., 2018).

### Cytokine profiles

Cytokine abundance in BAL was measured according to (Benedetto et al., 2019). Stimulatory potential of leukocytes was determined on freshly isolated MNC from blood and lungs, cultivated with distinct Toll-like receptor (TLR)-agonists or phorbol 12-myristate 13-acetate + ionomycin. Cytokines in the supernatant were determined by fluorescent assays (Ladinig et al., 2014) or a porcine TNF-α ELISA (R&D Systems).

### Peripheral blood mononuclear cell single-cell RNA sequencing

Cryopreserved PBMC were thawed and processed with Chromium Single Cell 3’ Library & Gel Bead v3.1 and Chromium Single Cell B Chip kits (10x Genomics, Pleasanton, CA, USA) to generate barcoded Gel Bead-In-Emulsions, according to manufacturers protocols. For each genotype and species constellation, equal numbers of cells from n=4 were pooled. Single cell libraries were prepared from 10,000 cells per pooled sample and sequenced with the NextSeq1000 (Illumina, San Diego, CA, USA) at a depth of > 30,000 paired reads per cell. Converted data were and aligned to Sscrofa11.1 (porcine) and GRCh37 (human), using Cell Ranger pipeline (v6.1.1, 10X Genomics). Quality check, dimensionality reduction, standard unsupervised clustering algorithms and the DEG analysis were performed using the standard Seurat pipeline (v4.3.0). Dimensionality reduction was done by PCA (20 components) and UMAP (resolution 0.8 for global clustering and 0.5 for monocyte sub-clustering). An adult human PBMC dataset (Bakken et al., 2021) was used as reference for data integration in Azimuth (https://azimuth.hubmapconsortium.org/).

### Phagocytosis and Oxidative Burst assays

Phagocytic potential of blood-derived monocytes and granulocytes was examined immediately after necropsy by Phagotest and Phagoburst kits (BD Biosciences). Phagocytosis was examined by stimulating blood samples with opsonized *E. coli* labelled with fluorescein isothiocyanate (FITC). Reactive oxygen species production was determined as the oxidation capacity of dihydrorhodamine to fluorogenic rhodamine. In humans, CD14^+^-cells were immunomagnetically isolated from PBMCs and incubated with fluorescently labelleled *Staphyloccocus aureus* and *Pseudomonas aeruginosa* bioparticles in an Incucyte SX1 life cell imagining incubator (Sartorius, Göttingen, Germany) over 48 hours, with automatic fluorescence MFI recording.

### Statistical analysis

Statistics for FC and IHC were performed in GraphPad Prism (GraphPad, La Jolla, CA) for calculating median and interquartile range (±). Two-tailed Wilcoxon matched sign ranked test assessed differences in paired analysis of littermates between CF and WT. Alternatively, two-tailed Mann-Whitney U test was performed for independent analysis. Significance threshold for Bonferroni corrected p-values was set at 0.05. For proteomics and scRNAseq, gProfiler (Raudvere et al., 2019) with Benjamini-Hochberg method for multiple testing adjustment was used, designating genes with adjusted P-values ≤ 0.05 and fold-changes ≥ 1.5 as significantly regulated. Functional enrichment analysis was done by ‘GO Biological Process’ ‘GO Molecular Function’, ‘GO Cellular Component’ and ‘KEGG’ databases.

For experimental details, antibodies, primers etc. see Supplementary Materials and Methods. Raw data on tissue weight, differential blood counts, qPCR, leukocyte and lymphocyte phenotypings, histological digital image analysis, cytokine profiles, differential monocyte receptors and phagocytic capacity are given in Table S11.

## Supporting information

Proteome profile of pig lung tissue

Proteome profile of pig pulmonary MNC

PBMC markers in pig and human scRNAseq

Monocyte markers in pig and human scRNAseq

Monocyte markers in human PBMC reference databases

Differential gene expression in cMO

Proteome profile of pig PBMC

## Acknowledgments

We are grateful to Maria Stadler (Institute of Immunology, University of Veterinary Medicine Vienna, Austria), Christian Eckardt and Tuna Güngor (Chair of Molecular Animal Breeding and Biotechnology, LMU Munich, Germany) for technical support.

## Author contributions

***F.J.*** *designed the study, performed tissue sampling, flow cytometry analysis, molecular analysis, interpretation and wrote the manuscript. He contributed to the breeding program and genotyping. **F.B.** performed tissue stainings and pathological analysis. **B.S., J.B.S.** & **T.F.** performed proteomic analysis. **G.S.** did bioinformatics analysis on scRNAseq data. **A. Sch.** characterized immune cells from patients. **S.Y.G.** & **M.A.M.** provided patient samples, made critical revisions and contributed to the writing of the manuscript. **A.B.** designed and led the breeding program of Cystic Fibrosis pigs. **M.C.-B.** performed qPCR analysis. **M.S.** contributed to qPCR analysis. **St.K. & H.B.** performed RNAseq experiments. **C.S.** performed PAEP assays. **D.Z.** contributed to scRNAseq experiments. **M.J.** contributed to the breeding program, genotyping and tissue sampling. **I.C.-P.** contributed to the phagocytosis assays**. K.K.** performed BAL analysis. **A.M.** & **K-L.L.** contributed to scRNAseq analysis and to the writing of the manuscript. **C.K.** contributed to interpreteation of scRNAseq. **E.W.** contributed to the design of the study and the writing of the manuscript. **L.M.** performed tissue stainings, pathological analysis and contributed to interpretation of the data. **W.G.** designed the study, performed flow cytometry analysis and led interpretation and wrote the manuscript **N.K.** designed and coordinated the study, led the interpretation and wrote the manuscript*.

## Data and materials availability

Raw data are available upon request.

