## Supplementary material for "Increased pulmonary monocyte infiltration and attenuated phagocytosis defines perinatal dysfunction of innate immunity in Cystic Fibrosis": Supplmentary Material

#### ***Table of content***

*page*


*Supplementary Methods* 2

*Supplementary Tables*

Table S1. Proteome profile of pig lung tissue *external Table S1.xls*
Table S2. Proteome profile of pig pulmonary MNC *external Table S2.xls*
Table S3. Clinical characteristics of patient cohort 10
Table S4. PBMC markers in pig and human scRNAseq *external Table S4.xls*
Table S5. Monocyte markers in pig and human scRNAseq *external Table S5.xls*
Table S6. Monocyte markers in literature 11
Table S7. Monocyte markers in human PBMC reference databases *external Table S7.xls*
Table S8. Differential gene expression in cMo *external Table S8.xls*
Table S9. Proteome profile of pig PBMC *external Table S9.xls*
Table S10. Transcriptional regulation of granule protein in monocytes 12

*Supplementary Figures*

Figure S1. Pathological assessment of CF specimens 13
Figure S2. Clustering of proteome profiles 14
Figure S3. Detailed examination of PAEP regulation 15
Figure S4. Gating strategies for major leukocyte subpopulations 16
Figure S5: Leukocyte subpopulations in distinct specimens 17
Figure S6. Gating for the respiratory Mononuclear Phagocyte (MP) Network 18
Figure S7. Cytokine profile from pulmonary mononuclear leukocytes 19
Figure S8. Cytokine profile from blood mononuclear leukocytes 19
Figure S9. RNA profiling of human and porcine PBMC 20
Figure S10. Analysis of phagocytic potential in Mononuclear Phagocytes (MPs) 21

*References* 22

**Experimental animals - production, genotyping and sampling**

With the offspring from previously generated *CFTR*$^{+/-}$ pigs (Klymiuk et al., 2012) a herd of heterozygous male and female pigs had been established. Animal production was performed under the supervision of the responsible regulatory authority (Regierung von Oberbayern) at the Center of Innovative Medical Models, LMU Munich. After 113 days of gestation, farrowing was introduced by administration of cloprostenol (Estrumate®, MSD Tiergesundheit by Merck, Kenilworth, NJ, USA). Rapid genotyping of newborn piglets was done within <6 hours with PCR detecting the WT allele (forward: *AGA AGA GTA GGG CCT TTG GCA T*; reverse: *AGC ACA TGT GGG TCT TAG AGT ACG*) or CF-KO allele (forward: *AGA AGA GTA GGG CCT TTG GCA T*; reverse: *TGG CTG AAC TGA GCG AAC AAG T*) (Ermund et al., 2018). Piglets were sacrificed according to the German Animal Welfare Act within the first day of life. Anesthesia was done with intramuscular injection of ketamine hydrochloride (Ursotamin®, Serumwerk Bernburg, Germany) and azaperone (Stresnil®, Elanco, Greenfield, IN, USA). Euthanasia was achieved by intracardial injection of embutramide, mebezonium and tetracain (T61®, MSD Tiergesundheit). Immediately, 20-30 ml of blood were collected by intracardiac puncture into heparinized tubes. For broncho-alveolar lavage (BAL), lungs were removed aseptically, sterile tubing was inserted into tracheae and the lungs were then washed with PBS (without $Ca^{2+}$ and $Mg^{2+}$, PAN-Biotech, Aidenbach, Germany). Washing was performed twice and lavages were pooled and placed on ice for cell counting. For mononuclear cell (MNC) isolation, lungs, including trachea, or entire spleens were explanted and placed in ice-cold PBS containing 1% Penicillin/Streptomycin (Thermo Fisher Scientific, Waltham, MA, USA). For other analyses, lung samples were taken from cranial, middle and caudal lung lobes. For histological assessment, fixation was done in 4% paraformaldehyde for 24-48 hours. For cryopreservation, samples were frozen on dry ice and cutted into small pieces (1x1x1 cm) and preserved at -80°C. Tissue was finally crushed with hammer & anvil and powderized by mortar & pestle, all under liquid nitrogen according to established procedures (Grotz et al., 2022) and stored at -80°C.

**Patient cohort – clinical characteristics and blood sampling**

The study was approved by the ethics committee of the Charité - Universitätsmedizin Berlin (EA2/016/18) and involved preschool children with previously diagnosed CF (3.9 ± 1.5 years) and age-matched healthy controls (3.2 ± 0.6 years) (each n=4). Healthy controls had no history of any chronic lung disease. Written informed consent to participate was provided by parents or legal guardians. Blood was drawn by peripheral venipuncture and collected in heparinized tubes.

**Isolation of mononuclear cells**

MNC were immediately isolated from blood, spleen or lungs under sterile conditions by gradient centrifugation from collected blood, spleen and/or lungs as reported previously (Rodríguez-Gómez et al., 2019). First, single cell suspensions were prepared for gradient centrifugation:

<u>Blood:</u> Heparinized blood was diluted 1:2 with PBS (20°C).

Spleen: Tissue was cut and filtered through a 70-µm cell strainer (Thermo Fisher Scientific), with cold PBS and a plunger from a sterile 10-ml syringe. The cell suspension was then centrifuged (10 min, 470 g, 4°C) and resuspended in 25 ml PBS (20°C) per sample.
Lung: Left and right lung lobes were chopped (3x4x3 mm) and rinsed with cold PBS to remove cells of blood-origin. Subsequently, lung tissue was digested in 50 ml RPMI 1640 with stable glutamine (PAN-Biotech) + 2% FCS (Sera Pro FBS, low endotoxin, PAN-Biotech) + 20 mM HEPES (Thermo Fisher Scientific) + 15,000 Units Collagenase type I (Thermo Fisher Scientific) + 1,250 Units Deoxyribonuclease I (Thermo Fisher Scientific) for 60 minutes at 37°C under permanent stirring. After digestion, the suspension was filtered through sterile metal sieves to remove remaining tissue. Next, dead cells were removed by filtration through sterile cotton wool. The resulting cell suspension was centrifuged (10 min, 420 g, 4°C) in 50-ml tubes and resuspended in 25 ml PBS (20°C) per sample.
Cell suspensions were subjected to gradient centrifugation with leukocyte separation medium adapted for newborn animals (Talker et al., 2013). Standardized leukocyte separation medium with 1.077 g/ml (PAN-Biotech) was diluted with PBS to 1.075 g/ml. 25 ml of cell suspension were layered above 15 ml of leukocyte separation medium in 50-ml tubes. After centrifugation (30 min, 900 g, 20°C), cells were collected from the designated leukocyte layer and washed twice with cold PBS. For proteomic profiling, pelleted cells were directly frozen in liquid nitrogen. Elsewise, cells were washed once more in cold RPMI1640 + 5% FCS, gently pelleted (10 min, 420 g, 4°C) and resuspended in 5 ml cold RPMI1640 + 10% FCS. Cells were then counted in Türk's solution (Merck) with help of a hemocytometer and cryopreserved in freezing medium in liquid nitrogen. (Leitner et al., 2012).

**Histopathological examination**

Fixed tissues of apical, cardiac, and caudal lung lobes were embedded in paraffin and cut into 2 µm sections. The lung sections were stained with HE or processed for immunohistochemistry. For the detection of $CD68^+$ cells (macrophages), lung sections were dewaxed and the epitope was retrieved by microwave heating (600W) for 15 min in 10 mM citric acid, pH 6.0 with 0.05% Triton X-100. The sections were incubated over night with a rabbit polyclonal anti-CD68 antibody (ab125212, abcam, Cambridge, UK; 1:1,000 dilution) followed by washing and incubation with a biotinylated secondary goat anti-rabbit antibody (BA-1000; Vector Laboratories, Newark, CA, USA; 1:200 dilution) for 1 hour. Antibody binding was visualized by incubation with Vectastain Elite ABC solution (Vector Laboratories), repeated washing and exposure to DAB (Merck). All sections were counterstained with hemalaun, dehydrated in ethanol, cleared in xylene, and coverslipped. Digital image analysis was based on standardized areas of 20x $2x10^6$ $\mu m^2$ per lung section with an adapted v9 nuclear count algorithm as described (Dietert et al., 2018).

**Mass spectrometry analysis**

*Sample preparation:* Lysis of cryo-powderized tissue, snap-frozen lung-derived MNC (isolated by gradient centrifugation) or PBMC was performed in 8 M urea/0.5 M $NH_4HCO_3$ with ultrasonication (18 cycles of 10 s) using a Sonopuls HD3200 (Bandelin, Berlin, Germany). Pierce 660 nm Protein Assay (Thermo Fisher Scientific) was used to measure and adjust protein concentration. Sequential digestion of 50 µg of protein was performed firstly with Lys-C

(FUJIFILM Wako Chemicals Europe GmbH, Neuss, Germany) for 4 h and subsequently with modified porcine trypsin (Promega, Madison, WI, USA) for 16 h at 37°C.

*Nano-liquid chromatography (LC)–tandem mass spectrometry (MS) analysis and bioinformatics:* MS, data acquisition and downstream data analysis were performed as described (Stirm et al., 2021). 1 µg of the protein digest was injected on an UltiMate 3000 nano-LC system coupled online to a Q Exactive HF-X instrument (both Thermo Fisher Scientific). Peptides were transferred to a PepMap 100 C18 trap column (100 µm×2 cm, 5 µM particles, Thermo Fisher Scientific) and separated on an analytical column (PepMap RSLC C18, 75 µm×50 cm, 2 µm particles, Thermo Fisher Scientific) at 250 nl/min flow-rate with a 160-min gradient of 3-25% of solvent B followed by 10-min increase to 40% and 5-min increase to 85%. Solvents A and B consisted of 0.1% formic acid in water and in acetonitrile, respectively. MS spectra were acquired using a top 15 data-dependent acquisition method on a Q Exactive HF-X mass spectrometer. MaxQuant (v.1.6.7.0) (Tyanova et al., 2016) software and the NCBI RefSeq Sus scrofa reference database (v.7-5-2020) were used for protein identification. All statistical analyses and visualization were performed using R (Team, 2021). Potential contaminants, identified by site and reverse hits were excluded from further analysis. Proteins having at least two peptides detected in at least three samples of each condition were quantified using the MS-EmpiRe algorithm (Ammar et al., 2019) as previously described (Flenkenthaler et al., 2021). Proteins with a Benjamini–Hochberg corrected P-value ≤ 0.05 and fold-change ≥ 1.5 were regarded as significantly altered. Principal component analysis and hierarchical clustering visualized as a heatmap (Gu et al., 2016) were used as unsupervised clustering methods. Over-representation analysis for significantly altered proteins was performed using WebGestalt (Liao et al., 2019) and the functional categories 'GO Biological Process nonRedundant' and 'KEGG'. Benjamini-Hochberg method was used for multiple testing adjustment. Top 15 pathways (based on significance) were used for visualization.

**Reverse transcription PCR**

RNA isolation from powdered lung tissue was done with Rneasy Microkit (Qiagen, Hilden, Germany). Approximately 100 mg of tissue were homogenized in 1 ml lysis buffer using a Polytron PT2500E (Kinematica, Luzern, Switzerland) in a pulsatile manner to avoid overheating. Further steps were carried out according to the Rneasy protocol. A total of 500-800 ng RNA was reversely transcribed by the SuperScript III kit (ThermoFisher Scientific). For *PAEP* splice variants, long-range RT-PCR was performed (Forward primer: GTTCAGGCCGTCGAAGTCAC; Reverse primer: GATTCCGCTGATGGGGAGGT), using HotStarTaq DNA polymerase (Qiagen) and *GAPDH* as housekeeping gene (Table M1).

qPCRs were run at 1:32 dilution of cDNA to avoid PCR inhibition. Stability of the house keeping genes *ACTB*, *GAPDH*, *TBP* and *YWHAZ* in neonatal lung tissue was compared by using RefFinder (http://blooge.cn/RefFinder/). In the following, the geometric mean of *ACTB* and *GAPDH* were used to normalize expression of genes of interest (Riedel et al., 2014). Comparative ΔCt analysis was performed using specific primers **(Table M1)** and FastStart Essential DNA Green Master (Roche Life Science, Basel, Switzerland) and Uracil-DNA glycosylase (Thermo Fisher Scientific). Optimized conditions were found at 2-step protocols, using a combined and extended step for annealing and polymerization. Assays were run on a LightCycler96 (Roche Life Science, Basel, Switzerland). The temperature profile was 50°C for

2 min, 95°C for 10 min, 45 cycles of 95°C for 10 s, 60°C or 63°C for 90 s. Data were analyzed on the LightCycler96 software (Roche Life Science, Basel, Switzerland), extracted and processed by Excel (Microsoft, Redmond, WA, USA).

***Table M1. Primer for quantification of mRNA levels in porcine lungs.***

| Gene name | Forward primer | Reverse primer | conc. (µM) | Annealing (°C) |
|---|---|---|---|---|
| ***ACTB*** | *AGCGCAAGTACTCCGTGTG* | *CGGACTCATCGTACTCCTGCTT* | 0.40 | 63°C |
| ***TBP*** | *GATGGACGTTCGGTTTAGG* | *AGCAGCACAGTACGAGCAA* | 0.55 | 63°C |
| ***YWHAZ*** | *GCATTATTAGCGTGCTGTCTT* | *ATGCAACCAACACATCCTATC* | 0.25 | 63°C |
| ***GAPDH*** | *CAGAACATCATCCCTGCTTC* | *GCTTCACCACCTTCTTGATG* | 0.40 | 60°C |
| ***CFTR*** | *AACCTGAACAAGTTTGATGAAG* | *AAGGCAAGTCCACAGAAGGC* | 0.25 | 60°C |
| ***PAEP_short*** | *CCACAATGAAAGAGCTCCAG* | *TCTCCTGAGGGATGGATTCT* | 0.25 | 63°C |
| ***IL1A*** | *AGCCCAGATCAGCAACATAC* | *CACTGCCTCATCCAGGTTATT* | 0.40 | 60°C |
| ***IL1B*** | *GTCCTCTGTCCTTGGCACC* | *GGCACACTCACCCCAAAGAA* | 0.25 | 63°C |
| ***IL18*** | *ACGATGAAGACCTGGAATCGG* | *AAACACGGCTTGATGTCCCT* | 0.55 | 63°C |
| ***TNFA*** | *CCCCTTGAGCATCAACCCTC* | *ATTGGCATACCCACTCTGCC* | 0.55 | 63°C |
| ***TGFB*** | *CCTGGGCTGGAAGTGGATTC* | *CCGGGTTGTGCTGGTTGTA* | 0.40 | 63°C |
| ***CYBB*** | *AGGCAGACTCAAGGCATTCAA* | *GCGCAGACCCAAGAAGTTTT* | 0.25 | 63°C |
| ***NLRP3*** | *AGACAGGAATGCACGTCTGG* | *TTGCATCTTGGCTGAGGTCC* | 0.25 | 63°C |

### Immunophenotyping

The cryopreserved MNC were thawed, counted and 1 x$10^6$ cells per panel were stained in round bottom 96-well micro-titer plates (Greiner Bio-One, Kremsmünster, Austria) as described (Rodríguez-Gómez et al., 2019). Flow Cytometry (FC) analysis was done, using established monoclonal antibody sources, labeling strategies and secondary reagents **(Table M2)**. Extracellular antigens were labeled in 3 incubation steps and free binding sites of secondary antibodies were blocked with anti-mouse IgG ChromePure™ (Jackson ImmunoResearch, West Grove, PA, USA). All incubation steps took place for 20 min at 4°C, each followed by two washing steps. A washing step comprised the addition of 200 µl staining buffer, centrifugation (4 min, 470 g, 4°C), discard of supernatant and the use of a plate shaker. After blocking, cells were washed with PBS (4°C), in order to prevent interference of the staining buffer with the fixable viability dye stain eFluor™ 780 (Thermo Fisher Scientific) of the third extracellular labelling. Thereafter, cells were fixed and permeabilized with the eBioscience™ Intracellular Fixation & Permeabilization Buffer Set (ThermoFisher). Staining of intracellular antigens, washing and final resuspension cells in 250 µl was performed with the provided buffer. Finally, cells were transferred to 5-ml tubes for flow cytometry analysis. Multicolor-stained leukocytes were analyzed with a FACSCanto II (BD Biosciences), equipped with three lasers (405, 488 and 633 nm). 5x$10^4$-2x$10^5$ leukocytes per sample, identified by their light scatter properties, were recorded by the FACSDiva™ software (BD Biosciences) and analyzed by FlowJo™ software.

***Table M2. Antibody panels used for FCM.***

| **Antigen** | **Clone** | **Isotype** | **Fluorochrom** | **Labeling strategy** | **Source** |
|---|---|---|---|---|---|
| ***Natural Killer cells*** | | | | | |
| CD3 | BB23-8E6- | IgG2a | PerCP-Cy5.5 | Directly conjugated | BD Biosciences |
| CD8α | 11/295/33 | IgG2a | PE | Two step biotin- | In-house[1] |
| NKp46 | VIV-KM1 | IgG1 | Alexa647 | Directly conjugated | In-house[2] |
| CD16 | G7 | IgG1 | FITC | Directly conjugated | BioRad |
| Ki-67 | B56 | IgG1 | BV421 | Directly conjugated | BD Biosciences |
| ***γδ T cells and cytotoxic T cells*** | | | | | |
| TCR-γδ | PPT16 | IgG2b | Alexa488 | Secondary | In-house |
| CD2 | MSA4 | IgG2a | PE-Cy7 | Secondary | In-house |
| CD27 | B30c7 | IgG1 | Alexa647 | Directly conjugated | In-house[2] |
| CD8β | PPT23 | IgG1 | PE | Two step biotin- | In-house[1,3] |
| Ki-67 | B56 | IgG1 | BV421 | Directly conjugated | BD Biosciences |
| ***B cells, regulatory T cells and activated & memory T cells*** | | | | | |
| CD4 | 74-12-4 | IgG2b | Alexa488 | Secondary | In-house |
| CD8α | 11/295/33 | IgG2a | PE-Cy7 | Secondary | In-house |
| CD25 | 3B2 | IgG1 | Alexa647 | Directly conjugated | In-house[2] |
| CD79α | HM57 | IgG1 | PE | Directly conjugated | Dako |
| Ki-67 | B56 | IgG1 | BV421 | Directly conjugated | BD Biosciences |
| ***Mononuclear phagocytes with expression of CD163 and CD80/86*** | | | | | |
| CD172a | 74-22-15A | IgG2b | Alexa488 | Secondary | In-house |
| CD163 | 2A10/11 | IgG1 | PE | Directly conjugated | BioRad |
| CD80/86 | CD152/Fc | Ig2b | PE-Cy7 | Secondary | Sigma-Aldrich |
| Ki-67 | B56 | IgG1 | BV421 | Directly conjugated | BD Biosciences |
| ***Mononuclear phagocytes with expression of CD11b and CD16*** | | | | | |
| CD11b | MIL4 | IgG1 | Alexa647 | Secondary | BioRad |
| CD16 | G7 | IgG1 | FITC | Directly conjugated | BioRad |
| CD163 | 2A10/11 | IgG1 | PE | Directly conjugated | BioRad |
| CD172a | 74-22-15A | IgG2b | PE-Cy7 | Secondary | In-house |
| Ki-67 | B56 | IgG1 | BV421 | Directly conjugated | BD Biosciences |
| ***Respiratory mononuclear phagocyte network*** | | | | | |
| MHC-II (SLA-DR) | MSA3 | IgG2a | PE-Cy7 | Secondary | Kingfisher |
| CD14 | CAM36A | IgG1 | BV605 | Secondary | BioRad |
| CD45 | K252.1E4 | IgG1 | Alexa647 | Directly conjugated | BioRad |
| CD163 | 2A10/11 | IgG1 | PE | Directly conjugated | BioRad |
| CD172a | 74-22-15A | IgG2b | Alexa488 | Secondary | In-house |
| Ki-67 | B56 | IgG1 | BV421 | Directly conjugated | BD Biosciences |

[a] anti-IgG2b-Alexa488 (Jackson ImmunoResearch)
[b] anti-IgG2a-PE-Cy7 (Southern Biotech, Birmingham, AL, USA)
[c] anti-IgG1-Alexa647 (Thermo Fisher Scientific)
[d] anti-IgG2b-biotin (Southern Biotech); Streptavidin - PE-Cy7 (Thermo Fisher Scientific)
[e] anti-IgG1-biotin (Southern Biotech); Streptavidin Brilliant Violet 605 (BioLegend, San Diego, CA, USA)
[1] biotinylation with EZ-Link™ Sulfo-NHS-LC-Biotin (Thermo Fisher Scientific)
[2] conjugation with Alexa Fluor® 647 Protein Labeling Kit (Thermo Fisher Scientific)
[3] non-biotinylated mAb commercially available from BioRad (Hercules, CA, USA)

**Cytokine profiles**

Cytokine levels of BAL were determined using Quantikine ELISA kits (R&D Systems, Minneapolis, MN, USA) as described (Benedetto et al., 2019). Freshly isolated MNC from blood and lung tissue were examined for cytokine production by various Toll-like receptor (TLR) ligands or phorbol 12-myristate 13-acetate (PMA) + ionomycin **(Table M3)**. Quadruplicates of $2x10^5$ cells in 200µl medium (RPMI 1640 + 10% FCS + 1% Penicillin/ Streptomycin) were cultivated in 96-well plates for 18 hours (37°C, 5% $CO_2$). Then, supernatants were pooled for each stimulus, cells removed (400g, 5 min, 4°C) and two aliquots per stimulus stored at -80°C for cytokine quantification. Supernatants were analyzed for IL-1β, IL-8, CCL2, IFN-α, IFN-γ, IL-10, IL-12 and IL-4 by a porcine multiplex fluorescent microsphere assay at the University Clinics for Swine (University of Veterinary Medicine, Vienna, Austria) as previously described (Ladinig et al., 2014). TNF-α concentrations were determined by the Porcine TNF-α DuoSet® ELISA and the DuoSet® Ancillary Reagent Kit 2 (R&D Systems), according to the manufacturer's instructions. Reactions was measured by color intensity at 450 nm by a Tecan Sunrise ELISA plate reader (Tecan, Maennedorf, Switzerland) and concentrations were calculated from duplicate wells by Magellan software (Tecan). For heatmap presentation, median of cytokine production in CF was referred to the median values in WT and transformed into $log_2$-fold change values.

***Table M3. Stimuli for measuring cytokine production of MNC***

| Stimulus | Source | PRR | Concentration | Reference |
|---|---|---|---|---|
| PMA / Ionomycin | 1 | independent | 5 ng/ml / 500ng/ml | a |
| Pam3Cys-SKKK | 2 | TLR-2/1 | 0.75 µg/ml | b |
| LPS *E. coli* O111:B4 | 3 | TLR-4 | 1 µg/ml | c |
| LPS_*P. aeruginosa* | 4 | TLR-4 | 1 µg/ml | b |
| Flagellin_*P. aeruginosa* | 3 | TLR-5 | 1 µg/ml | b |
| *ODN 2216* | 3 | TLR-9 | 5 µg/ml | c |
| *ODN 2243* | 3 | *ODN 2216* control | 5 µg/ml | --- |

1 Sigma-Aldrich by Merck, St. Louis, MI, USA
2 EMC Microcollections, Tuebingen, Germany
3 Invivogen, Toulouse, France
4 Thermo Fisher Scientific
a (Rodríguez-Gómez et al., 2019)
b (Braun et al., 2017)
c (Auray et al., 2016)

**Single-cell RNA sequencing (scRNAseq)**

For scRNAseq, human and porcine PBMCs were processed with Chromium Single Cell 3' Library & Gel Bead v3.1 and Chromium Single Cell B Chip kits (10x Genomics, Pleasanton, CA, USA) to generate barcoded Gel Bead-In-Emulsions, according to manufacturer's protocols. Briefly, cryo-conserved samples were thawed in a water bath at 37°C and cells were transferred into a 15-ml conical tube using a wide-bore pipette tip and slowly diluted with pre-warmed media (37°C, RPMI 1640 + 10% FCS). After centrifugation (5min, 300 g), supernatant was discarded and cells were resuspended in ice-cold media (4°C) for counting. Subsequently, $5x10^5$ cells of each individual were transferred into a new tube for pooling (McGinnis et al., 2021) according to species and genotype constellation (four donors per genotype and species). After centrifugation (5 min, 300 g), supernatant was discarded and cells were washed twice with ice cold 1xPBS + 0.04%BSA, resuspended in the same buffer and filtered through a 40 µm filter and placed on ice for final determination of cell concentration. Gel Bead-In-EMulsions (GEMs) and single-cell sequencing libraries were generated from 10,000 cells of each sample with the Chromium Single Cell 3' Library & Gel Bead Kit v3.1 and Chromium Single Cell B Chip Kit. Libraries were sequenced using The NextSeq1000 (Ilumina, San Diego USA, P2 flowcell kit with 100) with depth of > 30 000 pair reads per cell.

Data were provided in BLC format, converted to FASTQ format and aligned to Sscrofa11.1 (porcine) and GRCh37 (human). The Cell Ranger pipeline (v6.1.1, 10x Genomics) was used for barcode processing and generating the single-cell gene counting matrix. Samples were de-multiplexed to produce a pair of FASTQ files for each sample. Reads containing sequence information were aligned using the reference provided with Cell Ranger based on the GRCh37 and Sscrofa11.1 reference genomes and ENSEMBL gene annotations.

PCR duplicates collapsed to a single Unique Molecular Identifier (UMI) count when matching the same UMI, 10x barcode and gene. All the samples were aggregated using Cell Ranger without normalization and treated as a single dataset. The R statistical programming language (v4.3.0) was used for further analysis. Count data matrix was read into R and used to construct Seurat object (v4.3.0). The Seurat package was used to produce diagnostic quality control plots and thresholds for further filtering. Filtering was used to detect outliers and high numbers of mitochondrial transcripts. These pre-processed data were then analyzed to identify variable genes, which were used to perform PCA. Statistically significant principle components (PCs) were selected by PC elbow plots and used for UMAP analysis. For FindClusters, function resolution modularity optimization techniques (Louvain algorithm) were applied choosing 0.8 as parameter. For sub-clustered objects the resolution parameter was set to 0.5. Analysis of porcine dataset revealed one cluster of platelets with high mitochondrial content and low UMIs/gene and therefore removed from further analyses. After identifying major immune cell types we performed sub-clustering of monocytes (human: Cl.4 & Cl.13 pig Cl.0) using subset() function in Seurat pipeline. Two small subsets expressing B- or T-cell lineage marker such as Pax5 or CD3 in common were not considered for further analysis, as they seemingly represented doublets. Volcano plots of differentially expressed genes in classical monocytes were created by the web app VolcaNoseR (Goedhart and Luijsterburg, 2020).

Gene annotations in scRNAseq are taken from the Sscrofa11.1 (www.ensembl.org). Ensembl IDs that did not refer to an orthologous gene in human were replaced by gene names identified by BLATing porcine transcripts to the human reference genome. This includes the following: ENSSSCG00000029414=FCN1, ENSSSCG00000036618=FCGR3A, ENSSSCG00000026302=MKI67, ENSSSCG00000007978=HBA1, ENSSSCG00000031912=CLEC4D, SLA-DQB1=HLA-DQB1, SLA-DMA=HLA-DMA, SLA-DMB=HLA-DMB, ENSSSCG00000031242=TSPO, ENSSSCG00000028871=CSTA, ENSSSCG00000000694=GAPDH, ENSSSCG00000001456=HLA-DQA1, ENSSSCG00000001455=HLA-DRB1, ENSSSCG00000035224=ABI3, ENSSSCG00000026430=DDX3Y, ENSSSCG00000012178=EIF2S3Y, ENSSSCG00000033457=CCL15, ENSSSCG00000029630=TAF10, ENSSSCG00000039544=RPS24.

**Phagocytosis and ROS production**

Phagocytic capacity of pig blood cells was examined by Phagotest™ (BD Biosciences) according to the manufacturer's instructions. Oxidative burst of pig blood cells was explored by Phagoburst™ (BD Biosciences), according to manufacturer's instructions. Measurements for both assays were done with FACScan (BD Biosciences), aquiring $1x10^4$ leukocytes per sample. Analysis was done with CellQuest™ and FlowJo™ softwares.

In humans, $CD14^+$ were immunomagnetically isolated from PBMCs (Miltenyi Biotec, Bergisch-Gladbach, Germany) and seeded on a 96-well plate in a density of $5 \times 10^5$ in 100µl RPMI. Bioparticles of *Staphyloccocus aureus* and *Pseudomonas aeruginosa* were obtained pre-labelled with the fluorescent dye pHrodo red (Thermo Fisher Scientific) and added to the assay. Cells were incubated at 37°C and 5% CO2 in an Incucyte SX1 life cell imaging incubator (Sartorius, Göttingen, Germany). MFI was automatically measured for each well every hour for a period of 48h. To assess the oxidative burst capacity of human granulocytes, heparinised whole-blood duplicates were used. After lysis of red blood cells (BD Pharm Lyse, Beckton&Dickinson, Franklin Lakes, USA), cells were incubated with catalase (100U/ml; Sigma-Aldrich, St. Louis, USA) and Dihydrorhodamin (DHR, 10mg/ml, Thermo Fisher, Carlsbad, USA) substrate for five minutes at 37°C. While one tube served as negative control (the other was stimulated with PMA (8µg) for 30 minutes. Afterwards, the samples were run immediately on the flow cytometer (FACS Canto II, BD). The granulocytes were identified by their light scatter properties and the stimulation index was determined by calculating the ratio between the mean fluorescence of the stimulated cells and the mean fluorescence of the unstimulated cells.

### Supplementary Tables

*Table S3. Clinical characteristics of patient cohort.*

| Clinical Characteristic | | Healthy controls | CF patients |
|---|---|---|---|
| **Number of patients** | | 4 | 4 |
| Age (years) | Mean (± SD) | 3.2 (± 0.6) | 3.9 (± 1.5) |
| Sex (female) | n (%) | 2 (50%) | 3 (75%) |
| **Genotype** | | | |
| *F508del/F508del* | n (%) | --- | 1 (25%) |
| *F508del/MF* | | --- | 3 (75%) |
| Pancreatic insufficiency | n (%) | 0 (0%) | 4 (100%) |
| Body weight (kg) | Mean (± SD) | 14.4 (± 3.0) | 16.1 (± 3.2) |
| Body weight percentile | Mean (± SD) | 40.8 (± 29.4) | 45.0 (± 32.8) |
| Height (m) | Mean (± SD) | 96.0 (± 10.6) | 104.8 (± 13.7) |
| Height percentile | Mean (± SD) | 47.8 (± 38.3) | 56.0 (± 29.7) |
| BMI (kg/m²) | Mean (± SD) | 15.5 (± 0.8) | 14.9 (± 1.7) |
| BMI percentile | Mean (± SD) | 48.0 (± 22.7) | 32.8 (± 35.6) |

*Table S6. Monocyte markers in selected literature*

| | | **cMo** | **iMo** | **ncMo** |
|---|---|---|---|---|
| **Hamers 2019** | general | CD14++/CD16- | CD14++/CD16+ | CD14+/CD16+ |
| | human blood | CD163+, CD36+, CD64/FCGR1A, CD63 | **HLA-DR+**, CD11c+ | CD64-, CD163-, CD93-, CD82-, CD14- |
| **Duterte 2019** | human blood | CD55, Mac2/LGALS3, TLR4, CD261/TNFRSF10A, **CD114/CSF3R** CD35/CR1, CD11b/ITGAM, CD274/PDL1, CD89/FCAR, CD181/CXCR1, CD14+, CD16-, **S100A8**, **S100A9** | CD127/IL7R, CD215/IL15RA, CD105/ENG, LAP, CD73/NT5E, CD138/SDC1, CD94/KLRD1, CD167a/DDR1, CD179b/IGLL1, CD154/CD40L, P2RX7, BNIP3L | CD102/ICAM2, CD52, TLR2, CD88/C5AR1, **CD16/FCGR3A**, C3AR, CD15/CSF1R, CD85d/LILRB2, CCR10, CX3CR1, CD14-, SIGLEC10, SERPINA1, LILRA2 |
| **Olaloye 2021** | human intestine | n.a. | n.a. | n.a. |
| **Talker 2022** | general | CD14++/CD16- | CD14++/CD16+ | CD14-/CD16+ |
| | cattle PBMC | **CD62L/SELL**, IL1A, IL1B, IL1R, S100A7, **S100A8**, **S100A9**, **S100A12**, **ALOX5AP**, GSDME, NLRP3, VIM, CXCL2, CXCL4, CCR2, LYZ, VCAN, TGFBR1, TGFBI, IL12B, TNFSF8, TNSF14, CLECL1, GAPDH, ALDOA, PGK1, LDHA, LDHB, SDHA, SDHD, **CXCR2,** IL15RA, **TREM1** | CD86+, **MHCII+**, CD163+, CD11c+, CCL4, CFD, CASP4, ANXA3, ADA, CXCL16, PECAM1, CD52, CFD, CD1E, **BOLA-DRA** | CD5+, CD8a, CD205, CD163-, CD11c-, IL20RB, CCL3, CCL4, GBP5, CR2, TGFBR2, VASH1, IL11RA, CCL16, SLAMF6, CD274, IDO1, VSIG4, PFKM, ITGAL, C1QA, MS4A7, LY75, CLEC14A |
| **Yang 2017** | general | CD14++/CD16- | CD14++/CD16+ | CD14+/CD16++ |
| **Snodgrass 2022** | general | CD14+/CD16- | CD14+/CD16+ | CD14-/CD16+ |
| | human PBMC | IL8- | IL6, CRP, | IL8+, MPO- |
| **Herrera 2021** | pig PBMC | n.a. | n.a. | n.a. |
| **Thomas 2017** | general | CD14++/CD16- | CD14++/CD16++ | CD14+/CD16++ |
| | human blood | | **HLA-DR+**, CD11c+, | CD36-, CCR2-, |
| **Wong 2011** | general | CD14++/CD16- | CD14++/CD16+ | CD14+/CD16++ |
| | human blood | **S100A12, ALOX5AP**, **CLEC4D**, CCR2, IL1R2, **CSF3R**, ASGR2, **SELL**, CD163, CD99, CD9, CD36, LGALS2, EBI2, **S100A9**, **S100A8**, THBS1, PROK2, IL8, GPX1, ANPEP, IL1B, ITGA5, DYSF, NRG1, ASGR2, ALDH1A1, **SLC2A3,** EGR1 | MARCO, GPR35, NRIH3, GFRA2, FPRL2, SCD, ABPOBEC3A, CLEC10A, TGM2, **HLA-DRB3**, **HLA-DRA**, **CD74**, CD40, **HLA-DOA**, GFRA2, NKG7, PLAC8, E2F2, GBP4, PLVAP, TIMP1 | VMO1, SIGLEC10, ICAM4, IL21R, CD79b, CXCR1, SH2D1B, CXCR7, HES4, CEACAM1, RAB37, IL12RB1, CLEC2B, ADA, P2RX1, P2RY10, VASP, RHOC; PALM, SVIL, VAV2, LYN, VASP, VAV2, CBL, C2, C3, C1QA, CDKN1C, C1QB, CLEC4F, TAGLN, TCF7L2, CTSL, VMO1, IFITM3, **ABI3** |
| **Kapellos 2019** | general | CD14++/CD16- | CD14+/CD16+ | CD14+/CD16+ |
| | | CD36, CCR2, CD64, **CD62L/SELL,** CXCR1, **CXCR2** | LYZ, S100A8, **S100A10**, **HLA-DR**, **CD74**, IFI30, **HLA-DPB1**, CPV, CD86, CCR5, | CXCR1, XCR4, |
| **Boyette 2017** | general | CD14++/CD16- | CD14++/CD16+ | CD14+/CD16++ |
| **ours** | pig PBMC-scRNAseq | SERPINB1, RETN, **S100A9**, **CXCR2**, CSTA, CD52, **ALOX5AP**, LTF, **TREM1**, **S100A12**, SOD2, **CSF3R**, SLPI, IFITM3, CLEC2D, C4BPA, CD24, CPD, **CLEC4D**, LCN2, FBXO9, **SELL, S100A8,** CD9dim, **SLC2A3,** EGR1dim | CD163, FN1, MPEG1, YBX3, ARG1, CD204, EMP1, RPL4, CTSH, EPS8, EIF2A, RPS2, **SLA-DRB1**, LGMN, LMNA, CCL14, **S100A10**, APOE, **SLA-DQB1**, **SLA-DQA1**, ATP6, **CD74**, PLVAPdim, TIMP1dim | **CD16**, TMSB10, RPS27, PPIA, **ABI3**, SLP1, EGR1, S100A4, CD9, PRDX1, CSRP1, ADGRG1, LY6D, PRDX2, CIDEB, ATP5MC3, PLVAP, ITGA4, STAT1, TIMP1dim |

Color pattern according pig monocyte profiling. Red=consistent, yellow=partially consistent, blue=conflicting

*Table S10. Monocyte markers in selected literature*

| | human | | | | pig | | | | |
|---|---|---|---|---|---|---|---|---|---|
| | **WT** | | **CFTR** | | **WT** | | **CFTR** | | |
| **human_gene** | **avg_log2FC** | **p_val_adj** | **avg_log2FC** | **p_val_adj** | **avg_log2FC** | **p_val_adj** | **avg_log2FC** | **p_val_adj** | **pig alias** |
| *ELANE* | d.n.a. | | d.n.a. | | d.n.a. | | d.n.a. | | *ENSSSCG00000029130* |
| *PRTN3* | d.n.a. | | d.n.a. | | d.n.a. | | d.n.a. | | *ENSSSCG00000013417* |
| *CTSG* | d.n.a. | | d.n.a. | | d.n.a. | | d.n.a. | | *ENSSSCG00000037475* |
| *MPO* | d.n.a. | | d.n.a. | | d.n.a. | | d.n.a. | | *ENSSSCG00000017637* |
| *S100A8* | 1,05999162 | 1,50E-46 | 0,25597403 | 9,33E-21 | 0,77123422 | 4,97E-10 | 1,88795082 | 7,64E-102 | *S100A8* |
| *S100A9* | 0,64482205 | 5,00E-30 | 0,66097643 | 4,80E-36 | 0,94712724 | 1,11E-23 | 1,84560029 | 1,98E-111 | *S100A9* |
| *FCN1* | 0,31384522 | 2,15E-13 | d.n.a. | | d.n.a. | | 0,27602865 | 9,96E-20 | *ENSSSCG00000029414* |
| *LBP* | d.n.a. | | d.n.a. | | d.n.a. | | d.n.a. | | *ENSSSCG00000028758* |
| *LCN2* | d.n.a. | | d.n.a. | | 1,32954893 | 1,29E-15 | 1,49841898 | 2,25E-38 | *ENSSSCG00000005638* |
| *AZU1* | d.n.a. | | d.n.a. | | d.n.a. | | d.n.a. | | *AZU1* |
| *CD177* | d.n.a. | | d.n.a. | | d.n.a. | | d.n.a. | | *ENSSSCG00000003051* |

Values indicate differential regulation level (avg_log2FC) in classical monocytes of WT and CF PMBC pools. d.n.a … data not apparent in raw data set.

### Supplementary Figures

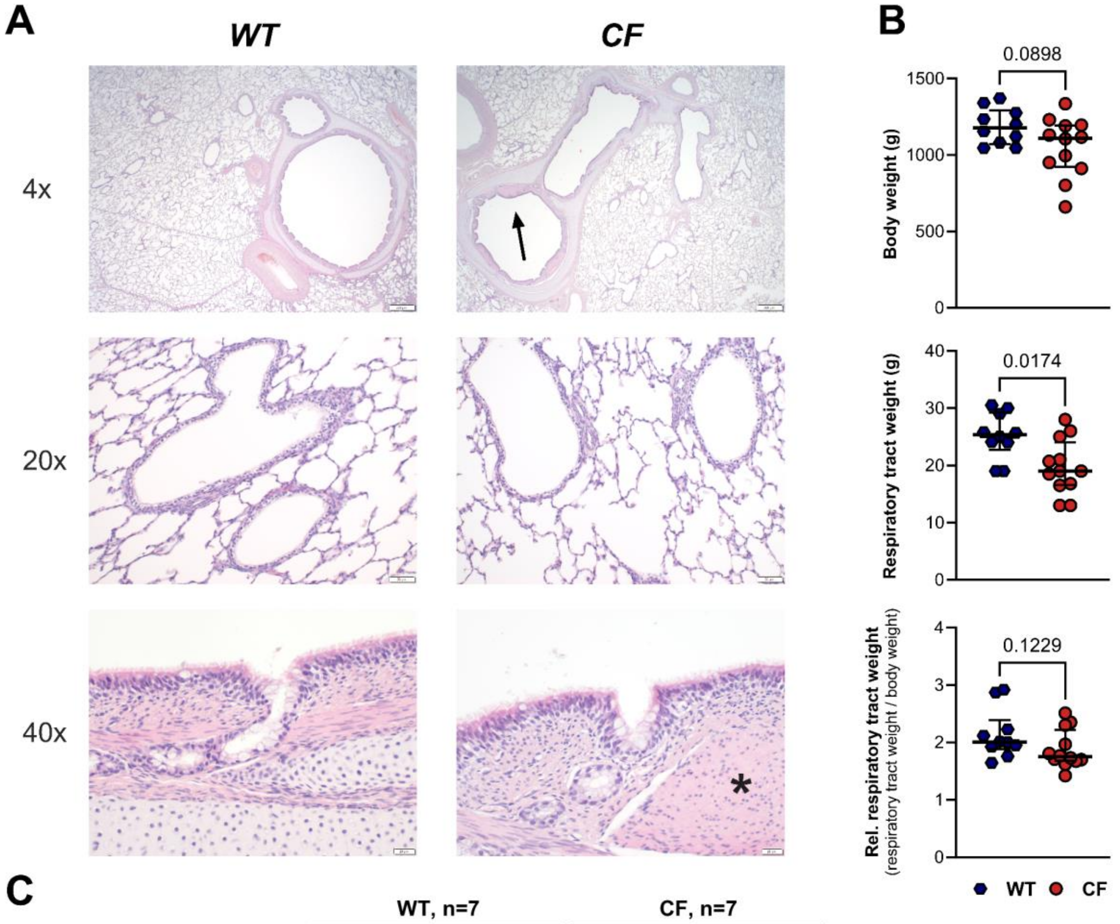


**C**

| Item | WT, n=7 Median | WT, n=7 IQR | CF, n=7 Median | CF, n=7 IQR | p-value |
|---|---|---|---|---|---|
| **WBC count, x10$^3$/µl** | **8.32** | 4.75 | **6.54** | 3.53 | **0.38** |
| **Neutrophils, %** | **65.6** | 17.2 | **65.00** | 13.1 | **0.90** |
| **Eosinophils, %** | **0.10** | 0.30 | **0.20** | 0.80 | **0.89** |
| **Basophils, %** | **0.20** | 0.10 | **0.30** | 0.10 | **0.16** |
| **Lymphocytes, %** | **29.50** | 10.10 | **29.20** | 10.10 | **0.48** |
| **Monocytes, %** | **4.60** | 1.20 | **5.10** | 2.80 | **1.00** |
| **NRBC, %** | **3.50** | 5.70 | **2.00** | 1.80 | **0.55** |
| **RBC count, x10$^3$/µl** | **4.76** | 3.93 | **5.82** | 0.72 | **0.54** |
| **Hemoglobin, g/dL** | **9.30** | 6.70 | **11.70** | 4.60 | **0.22** |
| **Hematocrit, %** | **30.50** | 12.20 | **37.30** | 4.60 | **0.26** |
| **MCV, fL** | **63.6** | 7.20 | **65.00** | 4.80 | **0.14** |
| **MCH, pg** | **19.50** | 2.60 | **20.20** | 2.60 | **0.10** |
| **MCHC, %** | **31.00** | 0.80 | **31.00** | 1.40 | **0.63** |
| **RDW-SD, fl** | **36.30** | 10.50 | **39.60** | 4.30 | **0.80** |
| **RDW-CV, %** | **17.10** | 2.40 | **16.50** | 1.70 | **0.78** |
| **Platelet count, x10$^3$/µl** | **400** | 210 | **327** | 65 | **0.36** |
| **P-LCR, %** | **41.00** | 21.20 | **46.00** | 15.30 | **0.16** |
| **PCT, %** | **0.47** | 0.15 | **0.39** | 0.08 | **0.40** |

***Figure S1. Pathological assessment of pig CF specimens.*** *(**A**) Representative WT and CF upper airway histologies at different magnifications, HE stain. Prominent smooth muscle bundles in larger CF bronchi indicated by arrow (top, 4x) and asterisk (bottom, 40x). (**B**) Body weight (upper panel), weight of distal respiratory tract, including trachea (middle panel) and the relative respiratory tract weight per body weight (lower panel). Each dot represents data of one individual animal. Median ± IQR and p-values are indicated, Mann-Whitney U test. (**C**) Differential blood cell count of newborn pigs (n=7 each). Median, IQR and p-values indicated, Mann-Whitney U test.*

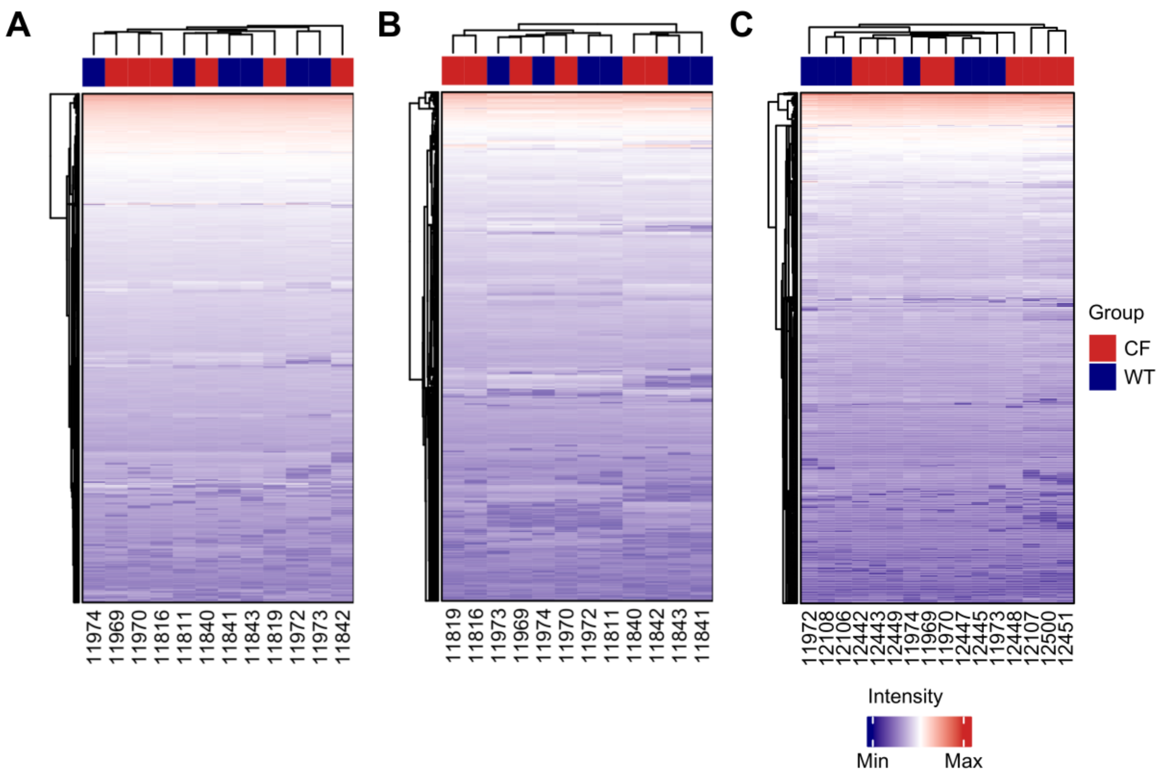


***Figure S2. Clustering of proteome profiles.*** *(**A**) Hierarchical clustering of log2 transformed label free quantification (LFQ) values from lung tissue samples, (**B**) purified pulmonary leukocytes and (**C**) peripheral blood mononuclear cells. (**D**) Principal component analysis of PBMC proteome profiles.*

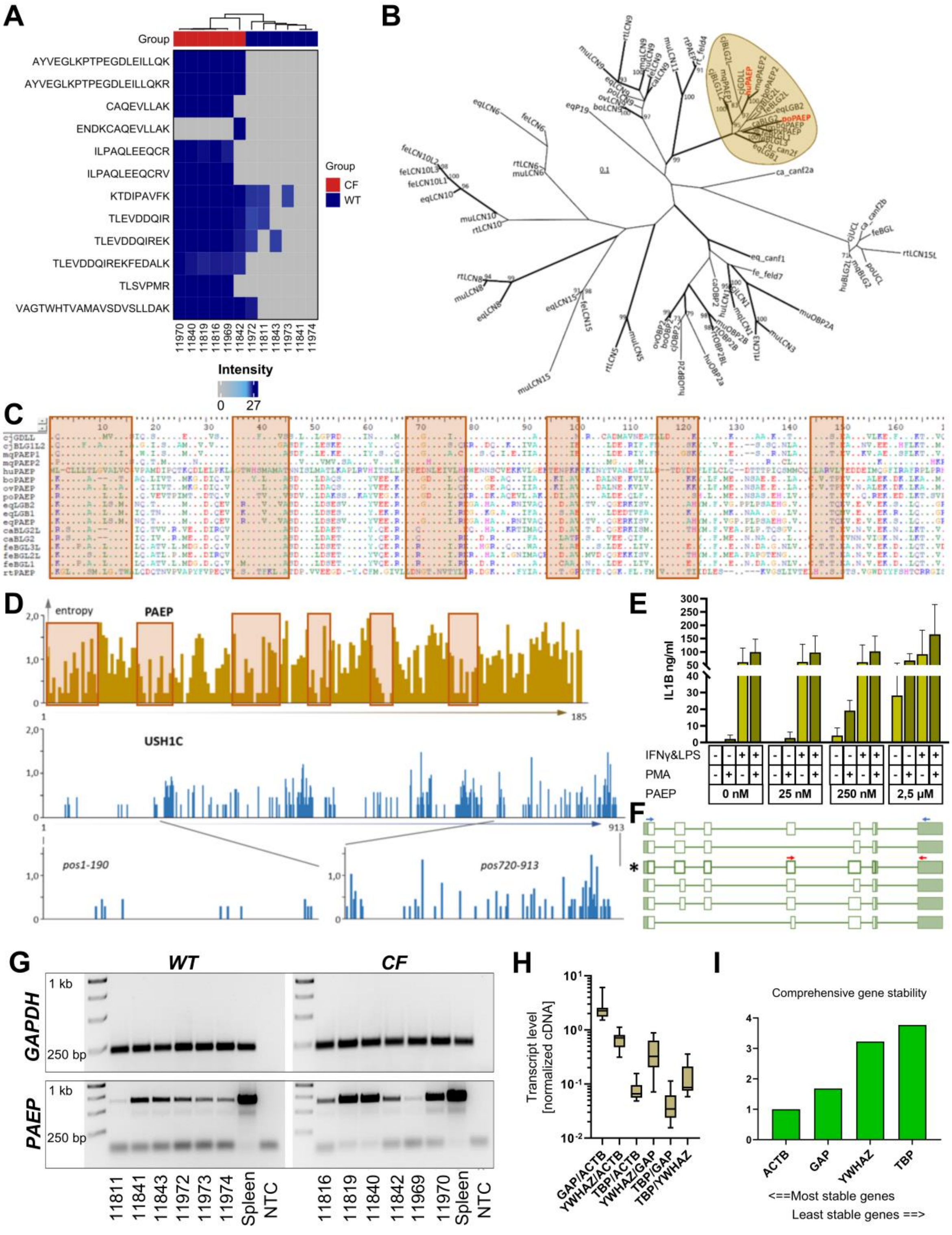


***Figure S3. Detailed examination of PAEP regulation.*** *(**A**) Proteomics revealed consistent abundance of multiple PAEP peptides in CF while this was lacking in WT. (**B**) Within proteins from the large lipocalin family, porcine PAEP (red) branches with proteins encoded by PAEP or BLG, LBG or GDLL genes in mammals (framed in gold), including the human PAEP (red). Protein sequences were extracted from annotated genes in the lipocalin loci of various mammalian species. Cj=marmoset; mq=macaque; hu=human; bo=cattle;ov=sheep; po=pig; eq=horse; ca=dog; fe=cat; rt=rat. In the PAEP cluster, human and pig encode a single PAEP gene, while other species constitute up to 3 genes. Phylogeneic tree constructed according to (Bartenschlager et al., 2022). (**C**) Proteins from the PAEP cluster appeared divergent, with only small regions indicating a higher degree of conservation (enboxed). (**D**) Entropy plots confirm low degree of conservation for PAEP proteins, compared to the designated "conserved" N-*

*terminal and "divergent" C-terminal regions of USH1C/Harmonin (see (Grotz et al., 2022)). (**E**) Indicated by IL-1$\beta$ production, human PAEP stimulated the human monocyte line THP1 after PMA conditioning in a dose-dependent manner. IL-1$\beta$ production induced by IFNG plus LPS was further enhanced by addition of high (2.5uM) PAEP concentrations. (**F**) Various splice variants have been predicted for human PAEP (according ncbi.nlm.nih.gov). Long range RT-PCR (blue arrows) amplified PAEP from lung tissue (**G**), confirming a single transcript variant, upon which primers for qPCR were designed (red arrows in (F)). (**H, I**) GAPDH and ACTB were identified as most stable house-keeping genes in neonatal lung for normalizing PAEP expression by RefFinder (http://blooge.cn/RefFinder/).*

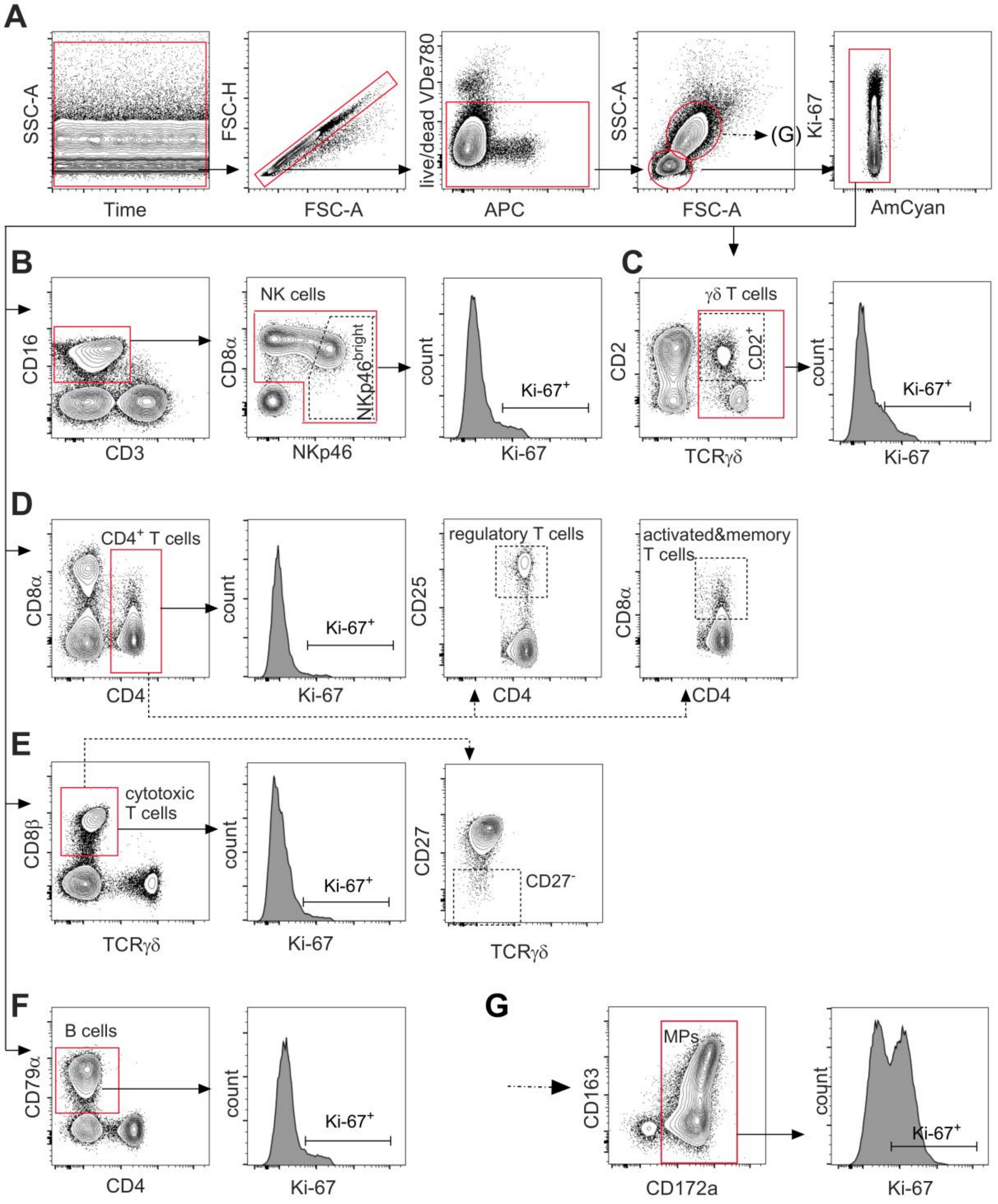


***Figure S4. Gating strategies for major leukocyte subpopulations.*** *(**A**) Identification of leukocytes by sequential gating using time parameter (to exclude potential clogging events), doublet discrimination and life/dead cell staining with VDeFluor780. Discrimination of lymphocytes and myeloid cells was performed based on FSC-A and*

*SSC-A light scatter properties. Lymphocytes were restricted to low AmCyan signals to exclude high auto-fluorescent artifacts and then gated for designated sub-populations by discriminating markers and their proliferation status by expression of Ki-67 (**B**-**F**). (**B**) NK cells were defined by a CD3⁻CD16⁺CD8α⁺NKp46⁺/⁻ phenotype. (**C**) γδ T cells were identified by TCRγδ expression and further discriminated by CD2. (**D**) CD4 T cells were identified by CD4 expression. CD4⁺CD25^high subpopulations were designated as regulatory T cells and CD4⁺CD8α⁺ cells as activated & memory T cells, respectively. (**E**) Cytotoxic T cells were characterized as CD8β⁺γδTCR⁻ cells and further discriminated by CD27 expression. (**F**) B cells were identified as CD4⁻CD79α⁺ cells. (**G**) Mononuclear phagocytes (MPs) were separated from lymphocytes by their FSC-A / SSC-A properties (A) and then defined as CD172a⁺ cells and examined for Ki-67 expression.*

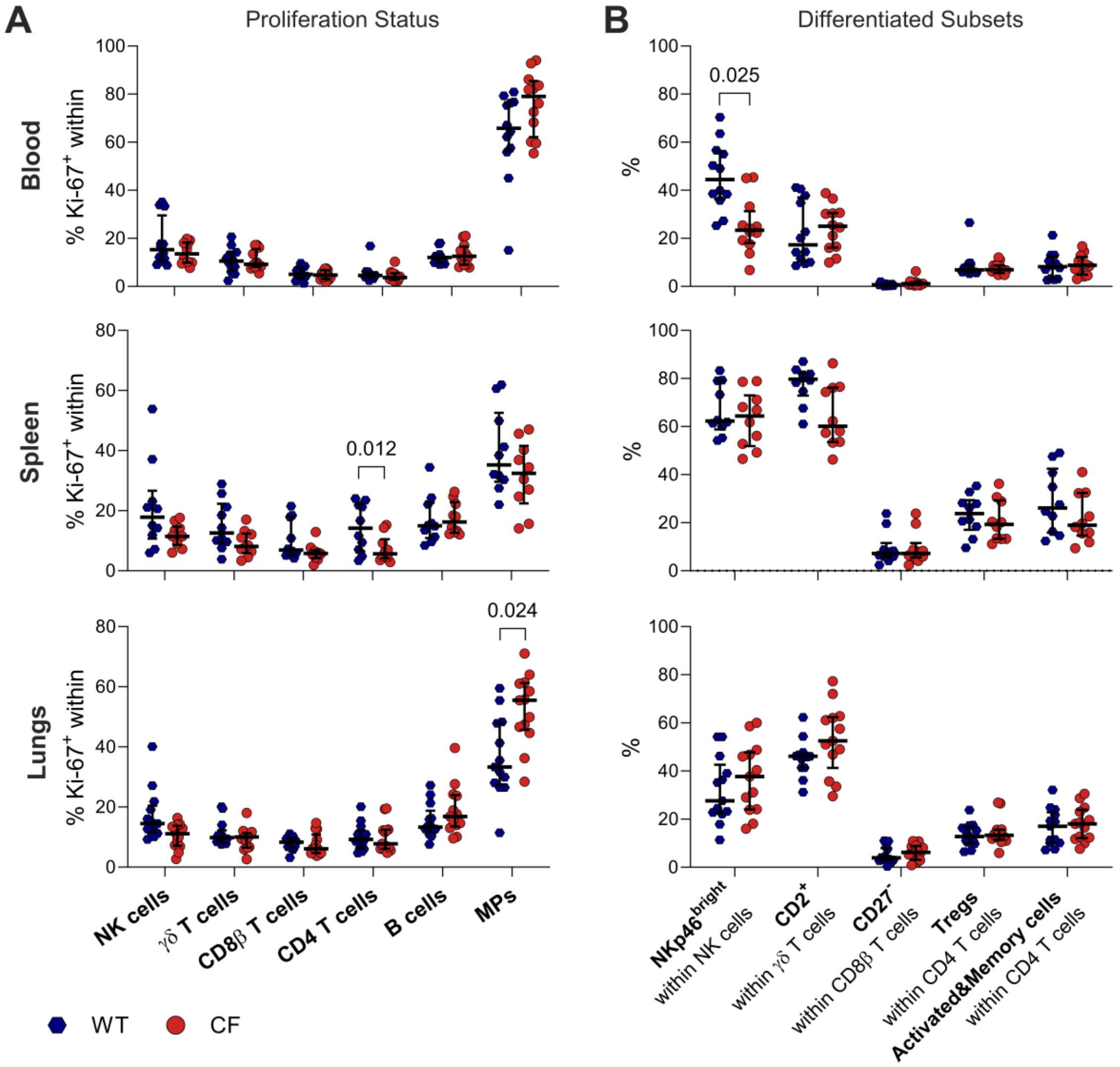


***Figure S5: Leukocyte subpopulations in different locations.*** *(**A**) Proliferation status within major leukocyte subpopulations from blood (n=12), spleen (n=10) and lungs (n=13) was assessed by Ki-67 expression. (**B**) Effector subsets in lymphocyte subpopulations in blood, spleen and lung, according gating strategies in Fig. S4. Each symbol represents data of one individual animal and median ± IQR are shown. Significant Bonferroni-adjusted p-values are indicated, Wilcoxon matched pairs signed-rank test.*

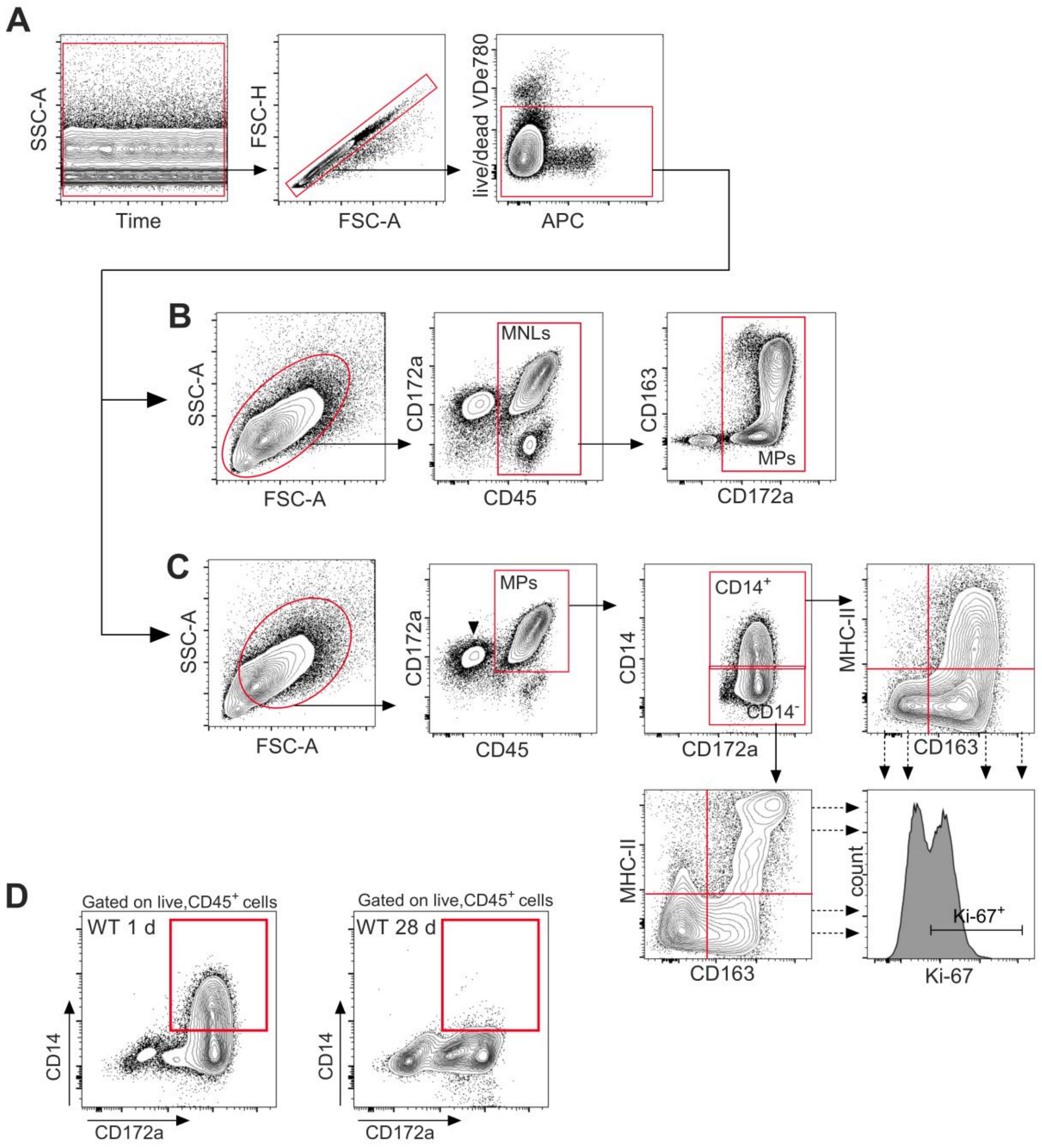


***Figure S6. Gating for the respiratory Mononuclear Phagocyte (MP) network.*** *(**A**) Sequential gating using time parameter (to exclude potential clogging events), doublet discrimination and life/dead cell staining with VDeFluor780. (**B**) Respiratory mononuclear leukocytes (MNLs) were identified according to light scatter properties and CD45 expression. MPs were identified by CD172a expression for determining the relative abundance of MPs within respiratory leukocytes. (**C**) Further characterization of MPs: here cells with low FSC/SSC signal (representing lymphocytes) were excluded; MPs defined by co-expression of CD172a and CD45 and further discriminated by CD14 expression. Further subtyping was based on CD163 and MHCII expression or the proliferation marker Ki-67. (**D**) Determining CD14$^+$ MPs after defining cut-off level in CD45$^+$ airway MPs from 28 day old WT pigs.*

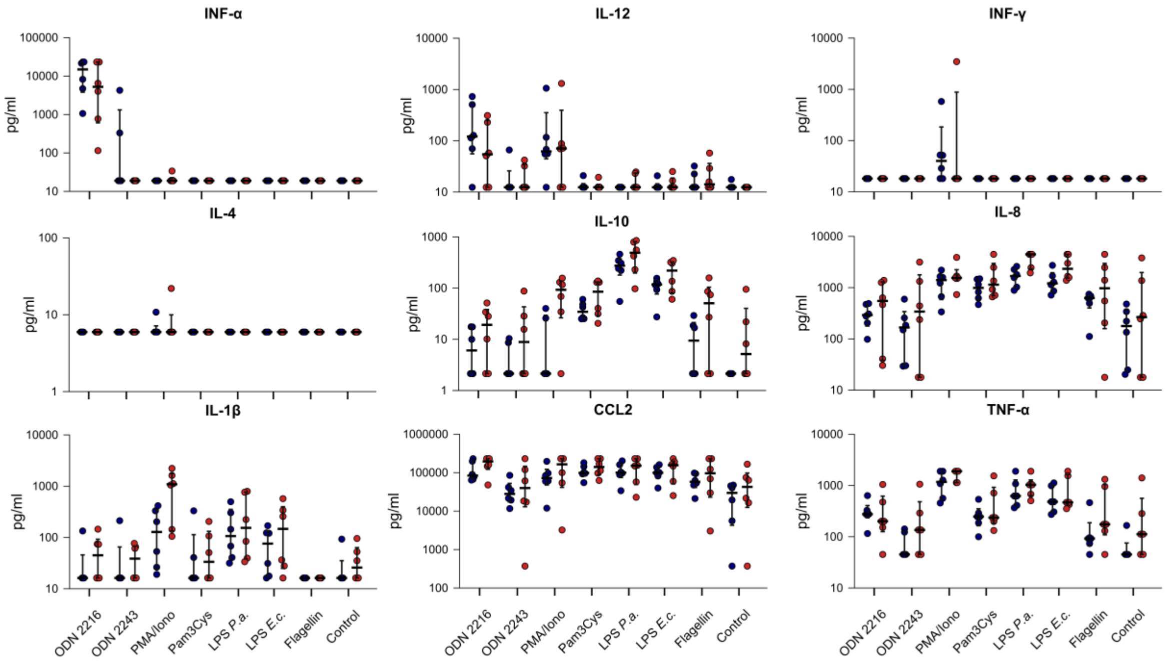


***Figure S7. Cytokine profile from pulmonary mononuclear leukocytes.*** *Freshly isolated mononuclear leukocytes from lungs of six littermate pairs of CFTR*$^{-/-}$*(red) and CFTR*$^{+/+}$*(blue) piglets were stimulated in vitro and supernatants were analyzed for cytokine production by multiplex fluorescent microsphere immunoassays. Each symbol represents data of one individual animal and median ± IQR are shown. Bonferroni corrected significance level of p<0.05 was not reached in any of the comparisons between genotypes, Mann Whitney U test.*

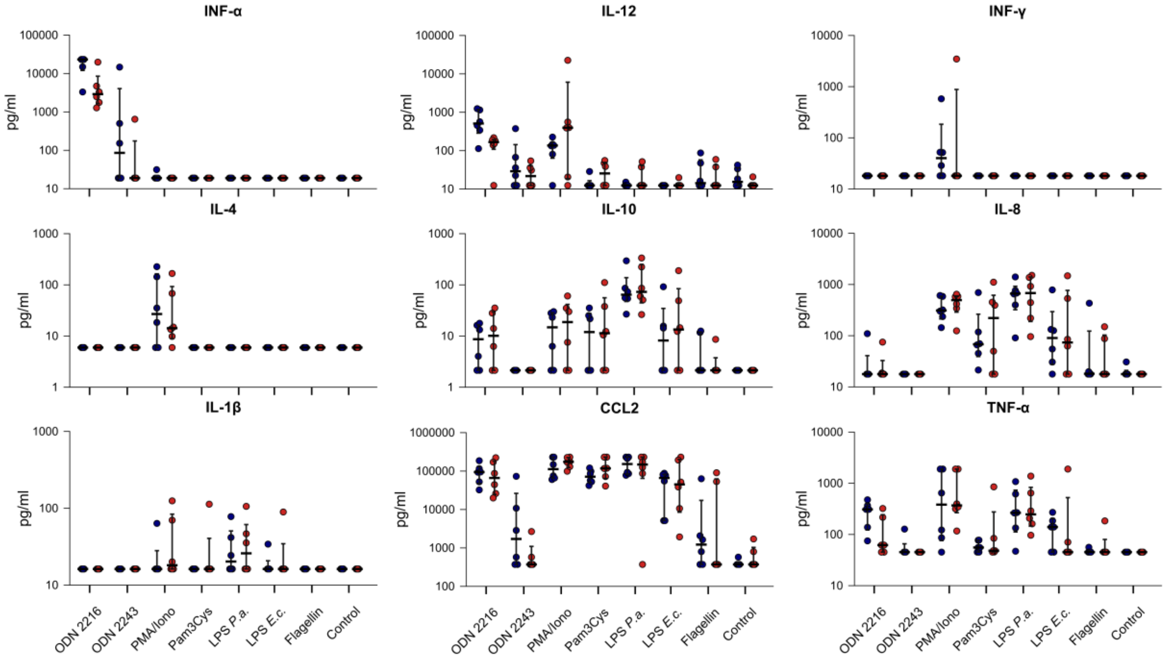


*Figure S8.* ***Cytokine profile from blood mononuclear leukocytes.*** *Freshly isolated PBMC from six littermate pairs of CF (red) and WT (blue) piglets were stimulated in vitro and supernatants were analyzed for cytokine production by multiplex fluorescent microsphere immunoassays. Each symbol represents data of one individual animal and median ± IQR are shown. Bonferroni corrected significance level of p<0.05 was not reached in any of the comparisons between genotypes, Mann Whitney U test.*

**A**

| Sample (PBMC) | Estimated Number of Cells | Fraction Reads in Cells | Mean Reads per Cell | Median Genes per Cell | Total Genes Detected | Median UMI Counts per Cell |
|---|---|---|---|---|---|---|
| Healthy Donor* | 11769 | 95.10 % | 54 286 | 1 906 | 23 036 | 6 521 |
| Children WT | 9 056 | 87.10 % | 57 086 | 1 842 | 22 042 | 5 990 |
| Children CF | 1 0979 | 84.00 % | 36 050 | 1 669 | 22 263 | 4 848 |
| Pig WT | 1 850 | 73.00 % | 297 878 | 1 672 | 14 193 | 5 381 |
| Pig CF | 2 582 | 71.00 % | 205 180 | 1 556 | 14 889 | 4 826 |

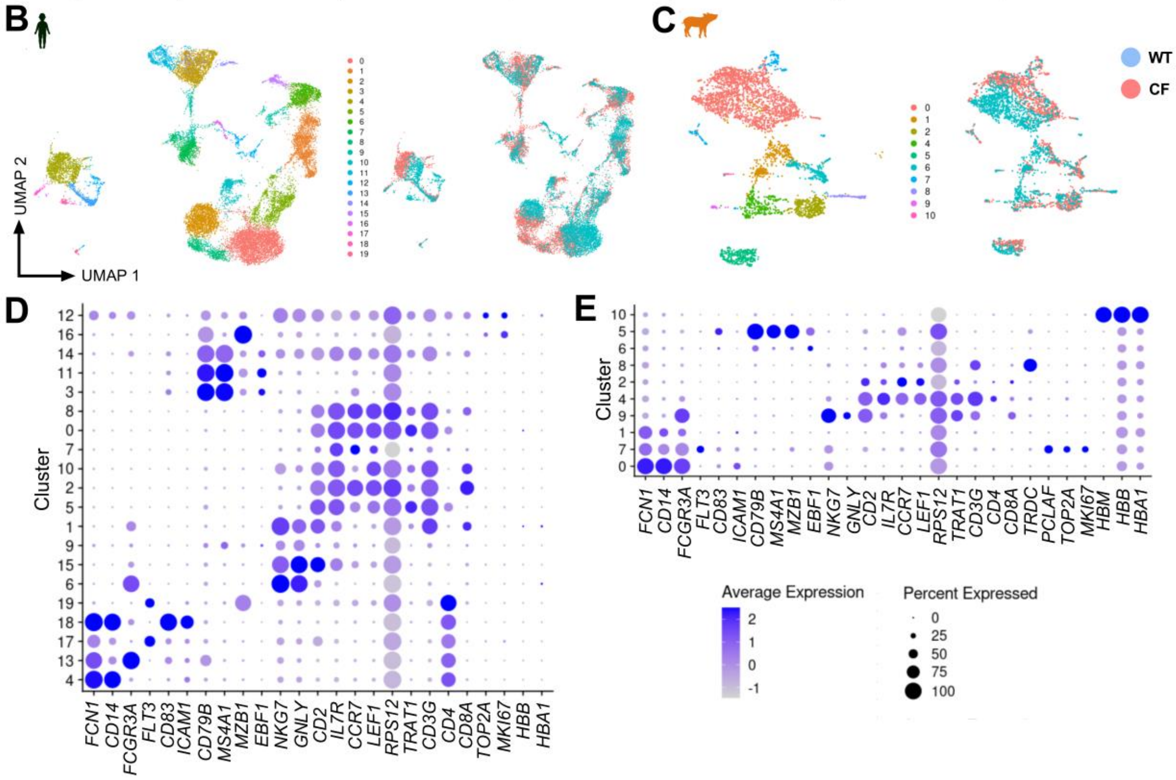


**F**

| Sample (PBMC) | Donors | B % (Count) | CD4 T % (Count) | CD8 T % (Count) | DC % (Count) | Mono % (Count) | NK % (Count) | Other T % (Count) | Other % (Count) |
|---|---|---|---|---|---|---|---|---|---|
| Healthy Donor* | 1 | **13.7** (1616) | **26.9** (3165) | **7.7** (907) | **2.1** (244) | **32.7** (3849) | **5.1** (602) | **7.1** (831) | **4.7** (555) |
| Children WT | 4 | **11.6** (1011) | **42.6** (3699) | **21.3** (1852) | **0.8** (72) | **10.8** (940) | **8.8** (766) | **3.2** (338) | **0.2** (15) |
| Children CF | 4 | **19.1** (2027) | **29.3** (3114) | **23.1** (2450) | **1.3** (140) | **10.1** (1077) | **11.7** (1246) | **5.0** (532) | **0.3** (34) |
| Pig WT | 4 | **7.7** (135) | **15.6** (275) | **0.9** (15) | **1.3** (23) | **59.6** (1047) | **1.0** (17) | **1.6** (29) | **12.3** (217) |
| Pig CF | 4 | **11.9** (263) | **24.9** (552) | **3.1** (69) | **4.5** (100) | **45.6** (1009) | **-** - | **1.4** (30) | **8.6** (191) |

**G**

| | B naive | B intermediate | B memory | Plasmablast | CD14 Mono | CD16 Mono | cDC | pDC | CD4 Naive | CD4 CTL | CD 4 Prol. | CD4 TCM | CD4 TEM |
|---|---|---|---|---|---|---|---|---|---|---|---|---|---|
| Children WT % (count) | 6.5 (562) | 2.8 (245) | 0.9 (82) | 1.4 (120) | 8.3 (718) | 2.6 (226) | 0.6 (49) | 0.3 (23) | 24.8 (2159) | 0.1 (6) | 0.5 (44) | 15.7 (1365) | 0.3 (24) |
| Children CF % (count) | 14.0 (1482) | 3.4 (364) | 1.3 135) | 0.4 (42) | 8.6 (918) | 1.5 (163) | 0.7 (79) | 0.6 (61) | 15.5 (1645) | 0.2 (21) | 0.1 (10) | 11.6 (1227) | 0.2 (20) |

| | Treg | CD8 NAIVE | CD8 Ptol. | CD8 TCM | CD8 TEM | dnT | gdT | ILC | MAIT | NK | NK Prol. | NK_CD56bright | HSPC | Platelet |
|---|---|---|---|---|---|---|---|---|---|---|---|---|---|---|
| Children WT % (count) | 1.1 (93) | 10.2 (8883) | 0 (3) | 0.6 (48) | 10.4 (906) | 0.8 (71) | 1.9 (162) | 0.1 (11) | 1.4 (121) | 7.8 (676) | 0.2 (17) | 0.8 (72) | 0.1 (5) | - (-) |
| Children CF % (count) | 1.2 (131) | 13.4 (1426) | 0.1 (9) | 0.4 (38) | 9.5 (1013) | 1.2 (125) | 2.7 (290) | 0.1 (14) | 1.3 (139) | 10.3 (1099) | 0.3 (27) | 1.2 (127) | 7 (0.1) | 0.1 (8) |

***Figure S9. scRNAseq profiling of human and porcine peripheral blood mononuclear cells.*** *PBMC from multiplexed human and pig of CF and control individuals (each n=4) were processed according to 10X Genomics protocols and analysed by using a human adult PBMC data set (https://support.10xgenomics.com/single-cell-gene-expression/datasets/3.0.0/pbmc_10k_v3) as reference.* ***(A)*** *CellRanger summary of scRNAseq QC parameters. Transcriptional heterogeneity of human* ***(B)*** *and porcine* ***(C)*** *PBMC depicted by UMAP dimensionality reduction. Left panels: UMAP clustering. Right panels: discrimination between CF (red) and WT (blue). Expression* *levels of selected transcripts in human **(D)** and porcine **(E)** UMAP clusters. Monocytes: FCN1, CD14, FCGR3A. Dendritic cells: FLT3, CD83, ICAM1. B cells: CD79B, MS4A1, MZB1, EBF1. NK cells: NKG7, GNLY. T cells: CD2, IL7R, CCR7, LEF1, RPS12, TRAT1, CD3G, CD4, CD8A. Proliferating cells: PCLAF, TOP2A, MKI67. Erythroid cells: HBM, HBB, HBA1. Dot sizes indicate the proportion of cells expressing the respective marker in a given cluster ("percent expressed"), colour-code identifies the expression level of transcripts ("average expression"). **(F)** Quantification of immune cell populations by using (https://azimuth.hubmapconsortium.org/) celltype.l1 annotations for human PBMC. **(G)** Quantification of human CF and WT datasets with celltype.l2 annotations.*

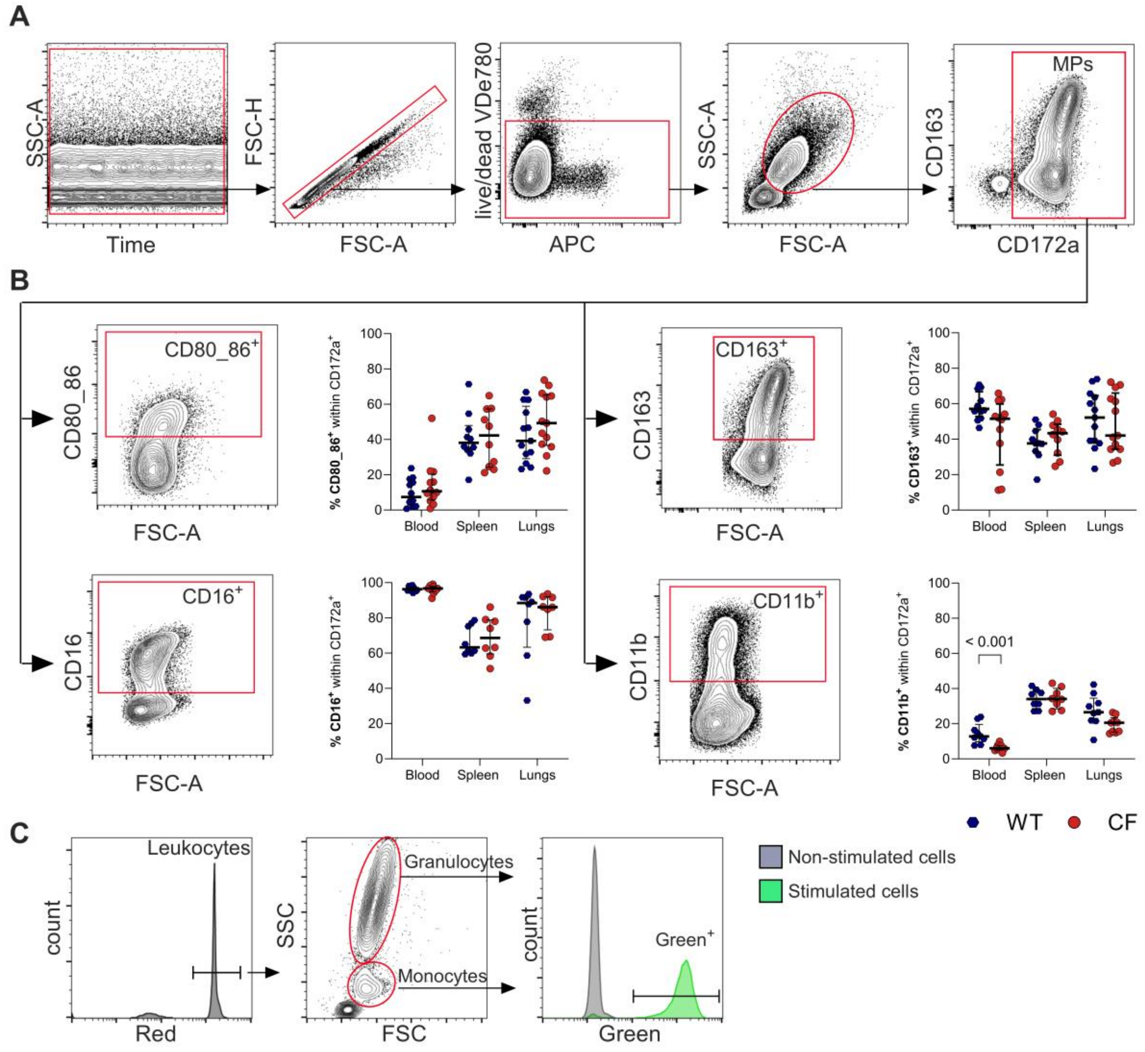


***Figure S10. Marker expression and phagocytic potential in Mononuclear Phagocytes (MPs).*** *(**A**) MPs were identified by sequential gating as described in Fig. S4A. Identification of CD80/86 **(B)**, and CD163 expression **(C)** (both: blood: n=12; spleen: n=10; lungs: n=13 per genotype) or CD16 **(D)** and CD11b expression **(E)** (both: blood: n=9; spleen: n=8; lungs n=8 per genotype). Left panels: Gates defining the respective markers. Right panels: proportions of CD80/86$^+$, CD163$^+$, CD16$^+$ and CD11b$^+$ sub-populations within CD172a$^+$ MPs in different locations. (**F**) Gating strategies for determining phagocytic potential in peripheral blood. Assay contained staining for DNA, facilitating the separation of leukocytes from debris or bacteria in the red channel. FSC and SSC parameters defined monocyte and granulocyte populations which were then assessed at 530/30nm (green), being indicative either for incorporation of FITC-labelled bacteria or the ROS-dependent production of rhodamine.*
